# The analysis of epigenomic evolution

**DOI:** 10.1101/2021.03.03.433796

**Authors:** Arne Sahm, Philipp Koch, Steve Horvath, Steve Hoffmann

## Abstract

While the investigation of the epigenome becomes increasingly important, still little is known about the long-term evolution of epigenetic marks and systematic investigation strategies are still withstanding. Here, we systematically demonstrate the transfer of classic phylogenetic methods such as maximum likelihood based on substitution models, parsimony, and distance-based to interval-scaled epigenetic data (available at [Github](https://github.com/Hoffmann-Lab/PhyloEpiGenomics)). Using a great apes blood data set, we demonstrate that DNA methylation is evolutionarily conserved at the level of individual CpGs in promotors, enhancers and genic regions. Our analysis also reveals that this epigenomic conservation is significantly correlated with its transcription factor binding density. Binding sites for transcription factors involved in neuron differentiation and components of AP-1 evolve at a significantly higher rate at methylation than at nucleotide level. Moreover, our models suggest an accelerated epigenomic evolution at binding sites of BRCA1, CBX2, and factors of the polycomb repressor 2 complex in humans. For most genomic regions, the methylation-based reconstruction of phylogenetic trees is at par with sequence-based reconstruction. Most strikingly, phylogenetic reconstruction using methylation rates in enhancer regions was ineffective independently of the chosen model. We identify a set of phylogenetically uninformative CpG sites enriching in enhancers controlling immune-related genes.

## Introduction

Sequence based methods for phylogenetic tree reconstruction developed more than half a century ago (Sokal and Michener 1958; Fitch and Margoliash 1967; Jukes and Cantor 1969; Fitch 1971), have laid the methodological foundation for much of the progress in evolutionary genetics. In addition to determining sequences of speciation events, they have allowed to associated genotypes with phenotypes and to investigate critical selection pressures (Pennacchio, et al. 2006; Kosiol, et al. 2008; Ge, et al. 2013; Gaya-Vidal and Alba 2014; Roux, et al. 2014; Reichwald, et al. 2015; Webb, et al. 2015; Sahm, et al. 2018; Cui, et al. 2019).

It becomes increasingly clear that the heritable information does not solely consist of the sequence of the four nucleobases adenine, cytosine, guanine and thymine (Boffelli and Martin 2012; Burggren 2016; Yi 2017; Lind and Spagopoulou 2018). While the functional analysis of chemical DNA modifications and its associated proteins is gaining pace, little is known about the long-term evolution and conservation of epigenomic signals. Among the most important of these epigenetic modifications are DNA methylation and post-translational modifications of histones (Chen, et al. 2017; Michalak, et al. 2019). DNA methylation marks, for instance, are copied to newly synthesized DNA strands by DNA methylation transferases targeted to the replication foci (Leonhardt, et al. 1992; Vertino, et al. 2002; Kar, et al. 2012). Importantly, epigenetic information may not only be passed on from cell to cell in the soma, but also through the germline from generation to generation (Verhoeven, et al. 2016; Perez and Lehner 2019).

However, it is not necessary to invoke these findings to take an interest at phylogenetic information conveyed by the epigenome. The evolution of epigenomic readers and writers themselves ultimately affects their function and changes in the epigenomic landscape may thus be understood as a consequence of this very process. While it is clear that the presence or absence of epigenetic marks in principle has a major influence on gene expression and cell identity, it is still largely open which marks have which functional significance where in the genome (Barrero, et al. 2010; Kim and Costello 2017; Xia, et al. 2020). As in many other examples (Bergmiller, et al. 2012; Luo, et al. 2015; Arun, et al. 2016), it is plausible that the degree of conservation would be a strong indicator for functional relevance. Furthermore, it could contribute to the elucidation of the molecular causes of phenotypic difference. To date, comparatively few studies have compared the epigenome of different species, e.g. to identify pairwise differentially methylated regions (Molaro, et al. 2011; Zeng, et al. 2012; Mendizabal, et al. 2016; Böck, et al. 2018)). To learn more about the long-term evolution of epigenomic marks, it appears necessary to develop new models allowing systematic evolutionary analyses -as is the case at the genetic level (Xiao, et al. 2014; Lowdon, et al. 2016). Obviously, the central question is whether the evolutionary information of individual marks is sufficient to allow meaningful analyses. Thus, it remains to be established which epigenomic marks are conserved well enough over longer evolutionary distances to allow the reconstruction of phylogenetic relationships.

That correct tree topologies can in principle be reconstructed from methylation data was shown by Martin et al. using blood samples from several primate species and a simple distance based tree method to reconstruct a single tree from the methylome (Martin, et al. 2011). Also, Qu et. al, whose main focus was on hypomethylated regions however, reconstructed a single tree from the whole methylome using a broader species set and a sophisticated time-continuous Markov chain model (Qu, et al. 2018). Their results indicated faster epigenomic evolution in rodent than in primate sperm. The work of Hernando-Herraez et al. was mainly concerned with differentially methylated regions resulting from pairwise comparisons, but also demonstrated, using a simple hierarchical clustering approach and great apes blood data, that genomic regions that showed incomplete lineage sorting on the nucleotide level recapitulated this on methylation level.

Building on these results, we systematically investigate different models of epigenomic evolution to facilitate insights into functions of single genes or pathways. Specifically, we transfer tree reconstruction such as Maximum Parsimony, Maximum Likelihood (based on substitution models), and distance-based methods to the level of DNA methylation. Subsequently, models are applied to simulated data as well as a publicly available real data to analyze their ability to reconstruct correct phylogenetic trees based on DNA methylation information. Substitution models arguably promise the greatest potential regarding functionally relevant analyses such as positive selection or accelerated epigenetic evolution in comparison to the genetic level. To this end, we are evaluating different evolutionary scenarios for different genomic features (e.g. enhancers, gene bodies).

## Materials and methods

### Real data set

To test the developed methods, we used publicly available whole-genome bisulfite sequencing data from blood samples of four primate species: *Homo sapiens* (hereafter, human), *Pan troglodytes* (chimpanzee), *Gorilla gorilla* (gorilla) and *Pongo abelii* (orangutan) (PRJNA286277,(Hernando-Herraez, et al. 2015)). The data set consisted of 286 to 324 million read pairs per species. With a length of 90 base pairs per read this amounts to 17-fold to 25-fold genome coverage, assuming a genome size of 3 giga base pairs per species (Table S1). Fig. S1 summarizes the processing of the real data, details are given in the supplementary material.

### Identification of orthologous defined regions and annotations

Our study distinguishes between four classes of genomic regions (Fig. 1A). The gene body class, reflecting the translated part of the genome, is differentiated from regulatory regions embedded in enhancer and two promoter-related regions reflecting potential differences in selection pressures of methylation in coding and non-coding sequences. Specifically, we consider promoter-near regions 2000 base pairs upstream and downstream the translation initiation site (TIS), respectively. In practice, the 2000-Up-TIS class covers the promotor and untranslated regions. Coordinates of the translation initiation and end sites of the human protein-coding genes (n=19,374) were obtained from the Uniprot track of the *University of California, Santa Cruz* (UCSC) table browser for the genome version hg38 (Karolchik, et al. 2004). Coordinates of enhancers were obtained from UCSCs geneHancer track retaining only “double elite” enhancers with known interactions (n=25,572). Coordinates were of 9,679 genes and 20,327 enhancers were successfully translated to the three non-human primates using the UCSC liftover tool (Kent, et al. 2010). For practical reasons, analysis was restricted to elements with a maximum length of 100,000/50,000 base pairs, respectively (Fig. 1A). For functional analyses, the UCSC hg38 *Transcription Factor ChIP-seq Clusters* track, aggregating transcription factor binding sites (TFBS) from more than 1,200 experiments in human samples for 340 transcription factors (TF) were incorporated into this study.

**Fig. 1.**
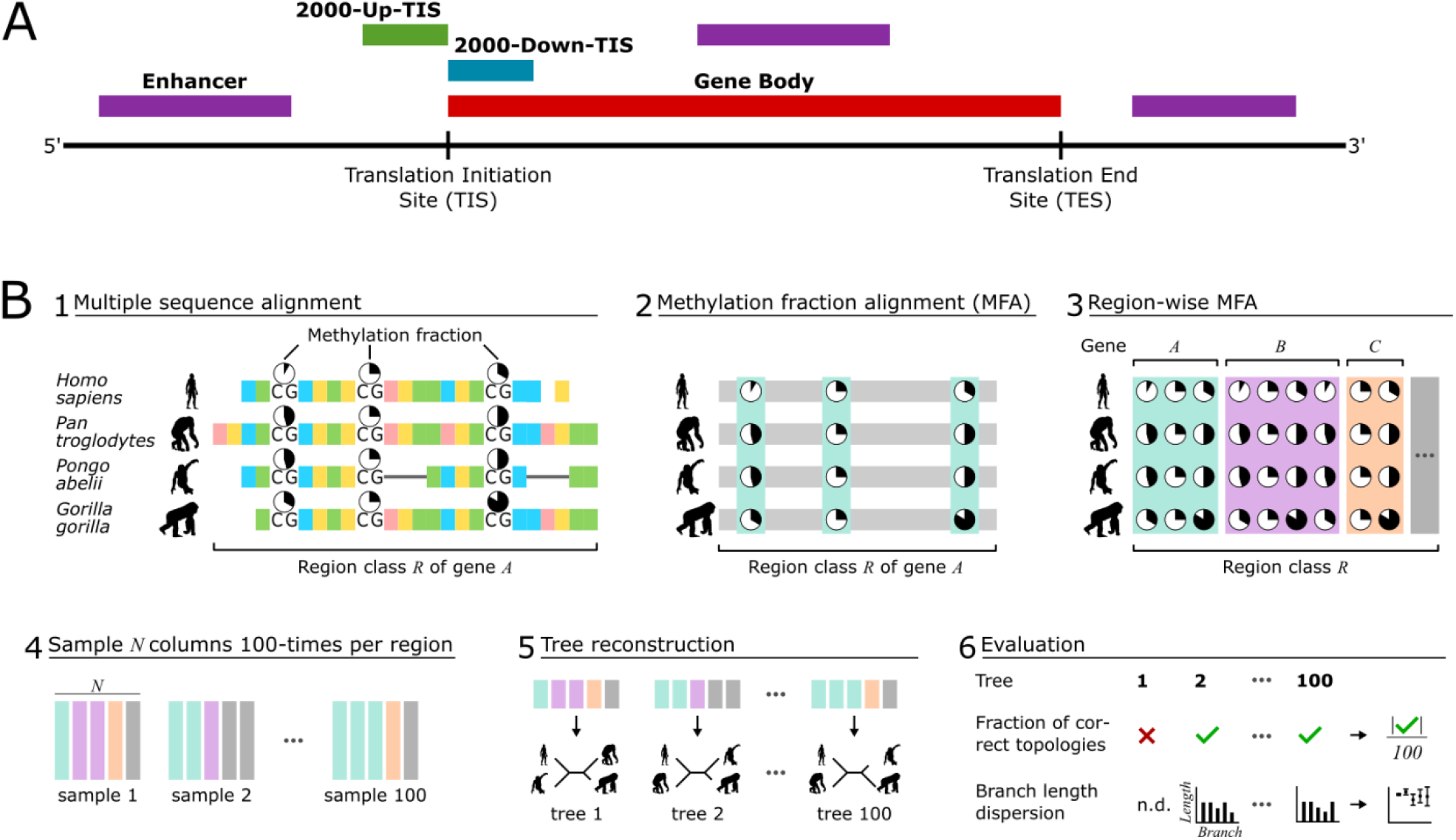
A) Classes of genomic regions examined in this work. The region classes were defined using the Uniprot annotation of all human protein-coding genes (2000-Up-TIS, 2000-Down-TIS, Gene body, n=19,374) or geneHancer annotation (Enhancer, n=25,572). The corresponding genome regions in chimpanzee, gorilla, and orangutan were acquired based on a genome alignment strategy. B) Workflow and evaluation strategy. i) For each region defined in A) of each gene a multiple sequence alignment of the four species studied in this work was created and the methylation fractions measured from blood samples mapped to the alignment. ii) From this, the methylation fraction alignment (MFA) was extracted. iii) We merged these gene-wise alignments to region-wise alignments. iv) From the region-wise methylation fraction alignments, we sampled 100 times *N* columns. v) From each of the 100 drawings, we reconstructed a phylogenetic tree. vi) As evaluation criteria, we used, on the one hand, the proportion of trees that correspond to the known great ape topology. On the other hand, we quantified the dispersions of the branch lengths. The procedure from iv) to vi) was performed for different values of *N* to consider the evaluation criteria as a function of the amount of input data.

### CpG alignment

For each region instance, the respective four orthologous sequences were aligned using Clustalw2, version 2.0.10 (Larkin, et al. 2007). Since the alignment of effectively non-homologous bases in poorly conserved regions may lead to false phylogenetic inference (Jordan and Goldman 2012), we used trimAl, version v1.4.rev15 with the parameter “-strictplus“ removing unreliable alignment columns (Capella-Gutiérrez, et al. 2009). In addition, only those alignments were considered for further analysis for which at least 25 evaluable CpGs remained. The methylation fractions previously determined in the individual species were then mapped to the CpGs of the alignment using the known coordinates. This led to methylation fraction alignments (MFA), forming the empirical data basis of this work (Fig. 1A).

### Evaluation strategy

The quality of individual tree reconstruction methods under different parametrizations was measured with increasing input sizes using two benchmarks: (i) the ability to correctly reconstruct the known primate tree topology and (ii) the degree of dispersion of the determined branch lengths. For these purposes, all methylation alignments of a region class were first concatenated (pooling). From this pool, *N* alignment columns were drawn for each combination of the used tree reconstruction approaches (see below) and *N* ∈ {100, 200,…,1000, 2000,…,10000}. The procedure was repeated 100 times. Subsequently, it was determined how many of the 100 repetitions led to the correct tree topology (Fig. 1B). To measure dispersion in terms of standard deviation of branch lengths from the mean length, the correct topology was fixed. Briefly, we expect to observe that the proportion of correct topologies should increase with increasing *N* and the dispersion of branch lengths should decrease.

### Tree reconstruction methods

We used three main strategies to reconstruct phylogenetic trees from the aligned methylation fractions: (i) Markov process based maximum likelihood, (ii) parsimony, and (ii) a distance-based methods. Here, we particularly focused on maximum likelihood and investigated different models, parameterizations, and assumptions based on this method. For simplicity, one combination is termed *default method* in the further and all alternative methods and parametrizations are compared against it. Fig. 2 summarizes the different tree reconstruction methods used. The developed methods for tree reconstruction from methylation (or more generally interval scaled) data were implemented in R and are available on [Github](https://github.com/Hoffmann-Lab/PhyloEpiGenomics).

**Fig. 2.**
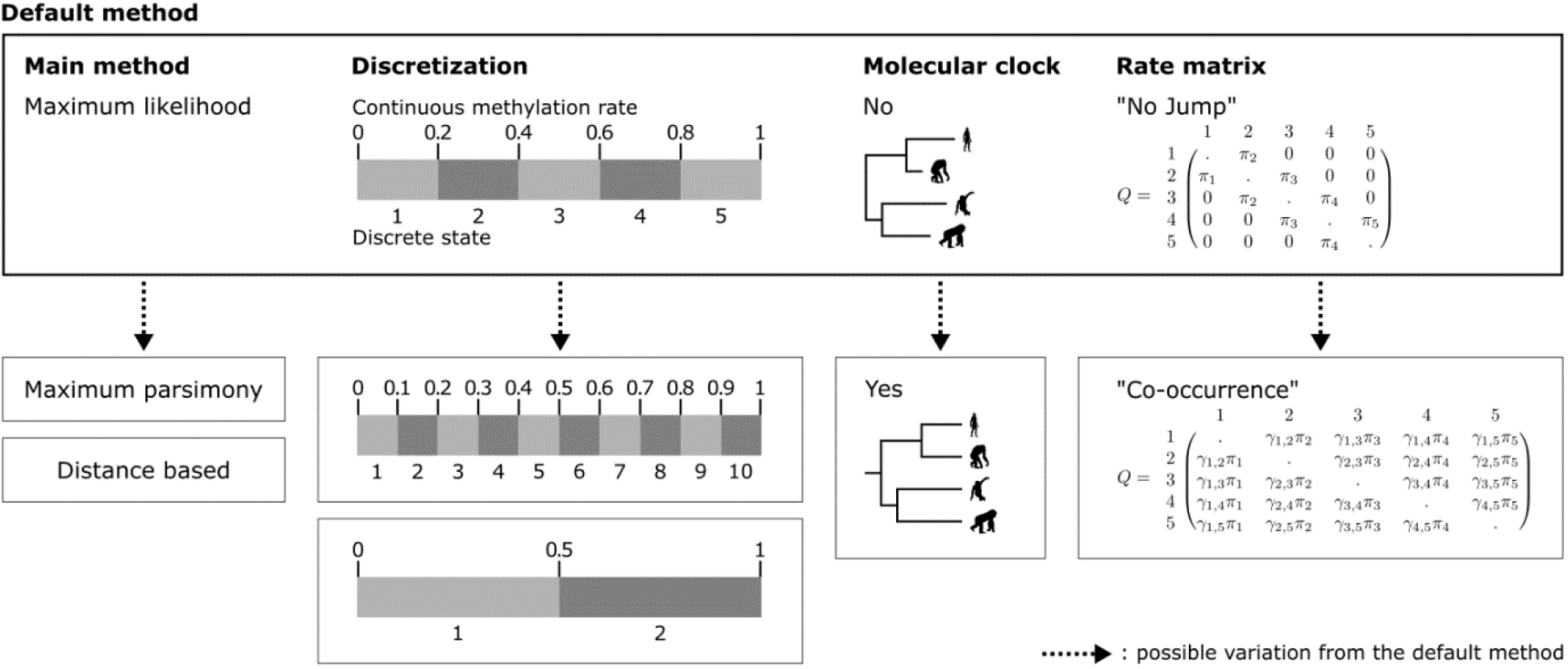
Applied tree reconstruction methods. Most of the analyses in this work were performed with a maximum likelihood tree reconstruction method based on a Markov model of the evolution of methylation fractions. The model is based on a discretization of the floating-point methylation fractions into 5 states. No molecular clock is assumed. The design of the transition rate matrix *Q* assumes that a methylation fraction status cannot evolve into a non-adjacent status within short time spans (*No Jump Model*). This means that when the methylation fraction changes from a state A to a non-adjacent state B during evolution, all states between A and B have been passed through. The combination of the tree reconstruction method and the described model properties is called the *default method* in the context of this work for simplification. To better assess the effects of the assumptions of the default method, we compared this method with variations of itself: different number of states, molecular clock or a transition rate matrix that allows the direct change to distant states. In addition, we have also applied two tree reconstruction methods that differ fundamentally from maximum likelihood. For this, either evolutionary distances based on the model of the default method were determined and then neighbor joining was applied or a parsimony approach was used.

### Maximum likelihood

We propose substitution models for the analysis of DNA methylation fractions analogous to the frequently used continuous-time Markov process models at nucleotide (e.g. (Jukes and Cantor 1969; Kimura 1980; Hasegawa, et al. 1985)), codon (e.g. (Goldman and Yang 1994)) and amino acid (e.g. (Kishino, et al. 1990)) level. In contrast to nucleotides, codons, and amino acids measured on a nominal scale, methylation fractions are measured on an interval scale. To address this difference, we introduce a discretization function

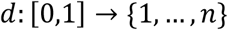

with *n* ∈ ℕ being the number of model methylation states and ∀ *x* > *y*: *d*(*x*) ≥ *d*(*y*).

Let a model *M* be defined by the pair *M* = (*π, Q*) consisting of an initial (and equilibrium) distribution *π* ∈[0,1]^*n*^and an *n x n*transition probability rate matrix *Q* = (*Q*_*s,t*_) with 1 ≤ *s* ≠ *t* ≤ *n*. As usual, the transition probability matrix *P*(*τ*) for a given branch length *τ* is determined by numerically finding a solution to

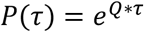

The likelihood of a given phylogenetic tree can then be determined by Felsenstein’s method assuming independent evolution of CpG sites (Felsenstein 1981). Branch lengths of a given tree topology are estimated by maximizing the likelihood and the optimal topology is that with the highest likelihood. For likelihood maximization we use an optimized version of the *Broyden*-*Fletcher-Goldfarb*-*Shanno* method (Broyden 1970; Byrd, et al. 1995).

In this paper, we propose two flavors of evolutionary models that we call *No Jump Model* and *Co-occurrence Model*. The No Jump Model assumes that the methylation fractions change smoothly during evolution, i.e. within short time intervals the methylation state of a CpG site can only change to one of the neighboring states (see Fig. 2). In contrast, the Co-occurrence Model also allows transitions to distant states in short time intervals. Here, the transition probabilities between two states are made dependent on how often these two states could be observed empirically within an alignment column.

To formally define these models, let *X* = (*X*_*i,j*_) be a given *k x l* matrix with the rows corresponding to *k* homologous CpG-sites, the columns corresponding to *l* indices of the examined species and each entry *X*_*i,j*_ ∈ [0,1] being the measured methylation fraction of the i^th^ CpG-site in the species with the index *j*;1 ≤ *i* ≤ *k*, 1 ≤ *j* ≤ *l*. Here, *X* will either be drawn from real data, i.e. the concatenated alignments of a region class, or from simulated data (see below).

Subsequently, the number of each methylation state *s* in each species *a* is counted using the function *o*_*a*_: {1,…, *n*} → ℕ_0_ with 1 ≤ *a* ≤ *l*

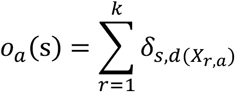

with *δ*_*p,q*_ being the Kronecker delta 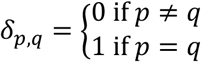

Based on this, we define the equilibrium frequency *π* = (*π*_*s*_) for both, the No Jump and the Co-occurrence Model as

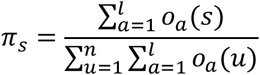

Accordingly, for the No Jump Model, we define *Q* = (*Q*_*s,t*_) as

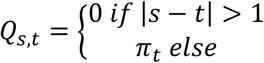

with 1 ≤ *s* ≠ *t* ≤ *n*. For the Co-occurrence Model, the number of co-occurrences of all combinations of methylation states s and t and species *a* and *b* is counted using a function *co*_*a,b*_: {1,…, *n*}^2^ → ℕ_0_ with 1 ≤ *a* ≤ *l*, 1 ≤ *b* ≤ *l, a* ≠ *b*

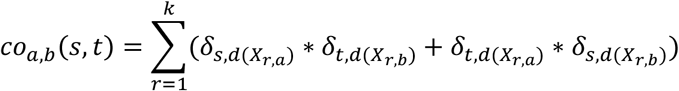

Let, 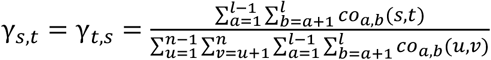 be the fraction observed co-occurrences of the states *s* and *t* across the alignment.

Finally, we define *Q* = (*Q*_*s,t*_) for the Co-occurrence Model as

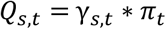

with 1 ≤ *s* ≠ *t* ≤ *n*. As usual, for both models *Q*_*s,s*_ are fixed such that

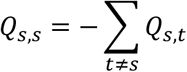

For this study, we tested the models with different parameters and technical settings. Detailed parametrizations for the No Jump and Co-occurrence models are shown in Figures S2 and S3. Specifically, we tested models with different state numbers *n*, and we used the models both with and without the assumption of a molecular clock.

### Maximum Parsimony

The basic idea of Fitch’s Maximum Parsimony algorithm (Fitch 1971) is to find a set of sequence states at the inner nodes minimizing the number of necessary changes along the edges of the tree. We adapted the algorithm to work on the interval scale. Figures S4-S6 illustrates the differences between the algorithm and our modification.

In the bottom-up post-order tree traversal, an interval *I* (*m*) is assigned to each node *m*. Leaves are initialized with

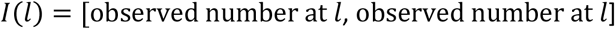

Then, for each inner node *m* with children *x* and *y, I* (*m*) is determined by

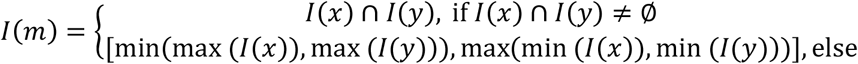

In the top-down pre-order tree traversal, each node *m* is assigned a number *i*(*m*) ∈ *I* (*m*). For the root *r, i* (*r*) is chosen as an arbitrary number within *I*(*r*); for all other inner nodes with parent node state *p*

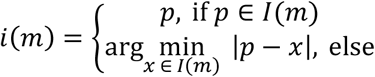

As with the original algorithm, the optimal tree topology is the one that explains the sequence alignment with the lowest number of necessary changes overall.

### Distance-based methods

Distance matrices were determined based on the *No Jump Model* described above. Each distance matrix

*T* = (*T*_*a,b*_) with 1 ≤ *a* ≤ *l*, 1 ≤ *b* ≤ *l* was determined by maximizing the likelihood for the pairwise distances in the usual way

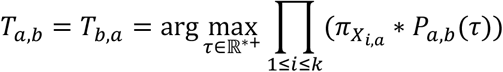

for *a* ≠ *b*; and *T*_*a,a*_ = 0 for 1 ≤ *a* ≤ *l*. For each distance matrix *T* the corresponding phylogenetic tree was reconstructed using the Neighbor Joining algorithm (Saitou and Nei 1987).

### Simulations

Artificial alignments were generated based on the known tree topology and divergence times of the great apes (Locke, et al. 2011). For each artificial alignment column, the tree was traversed in pre-order. For each node *m*, a state was drawn from (1,…, *n*) using a probability vector *φ*(*m*). For the root *r, φ*(*r*) was set to the equilibrium distribution *φ*(*r*) = *π*. For every other node *m* with the incoming edge *e* with length *τ*_*e*_ and the parent node in state *s, φ*(*m*) was set to the *s*-th row of the matrix *P*(*τ*_*e*_) (Fig. S7). The simulated alignment column then results from the states assigned to the leaves of this tree. This simulation approach has been widely used at nucleotide and codon level, e.g.(Rambaut and Grassly 1997; Yang 1997; Fletcher and Yang 2009).

This general scheme has been extended for the different concrete analyses. For the comparison of real and simulated data, noise of different orders of magnitude was added to the simulated data. For the analyses of long-branch attraction and resolution limits for reconstruction depending on the branch length the tree used for the simulation was changed. (see Supplement methods for details, also for the analysis of site-specificity that was conducted on real data).

## Results and Discussion

On nucleotide data, maximum likelihood based Markov models are critical tools for formal testing of evolutionary hypothesis. This includes the detection of sequences under positive selection on certain branches of a phylogeny. In order to address such questions in the epigenome, we first focused on the evaluation of maximum likelihood reconstructions in this work. Specifically, we investigated different flavours of this methodology. A discretization of CpGs according to their methylation levels is at the heart of the default method and derivatives thereof. Depending on the methylation, in our models, a single CpG can assume on of two, five (default method), or ten states.

### CpG Methylation varies widely between regions but little between species

To test the functionality of the developed methods, we used a public whole-genome bisulfite sequencing data set generated from blood samples of four great apes: human, chimpanzee, gorilla, and orangutan (Hernando-Herraez, et al. 2015) as well as to simulated data. On a genome-wide level, methylation patterns are extremely similar in all species and apparently dominated by the functional aspects of the four selected classes of regions (see Fig. 1A for region definition): In the gene body, the distribution of methylation fractions assume a shape resembling a beta distribution with prominent peaks at both ends of the unit interval. The peak for high to complete methylation rates is substantially higher than the peak for low to missing methylation (Fig. 3A). As methylation rates in the gene body positively correlate with gene expression, this observation may be explained with the transcriptional activity of the genes (Zemach, et al. 2010; Yang, et al. 2014). However, more than a quarter of the CpGs in the data set still show intermediate methylation between 0.2-0.8. After discretization, these levels correspond to the three intermediate states in the five-states model of the default method (Fig. 2). The methylation landscape of gene bodies is characterized by a sharp increase of average CpG methylation at the 5’-end and remains relatively stable at about 80% from this point on (Fig. 3B). On average, the gene body shows the highest methylation levels amongst all compared regions, but at the same time lowest CpG density by far: apart from small areas in vicinity to the 5’ and 3’ end, it is stable below 0.5% (Fig. 3B). For comparison: the genome-wide CpG fraction is 1% (Stevens, et al. 2013). The gene body also shows the lowest pair-wise methylation rate similarity among species (Table 1). Given the above-mentioned dependency of methylation and expression levels, this finding may reflect individual short-term expression changes, e.g. in the context of diurnal rhythms. In line with a characteristic hypomethylation in promotor regions, the distribution in the 2000-Up-TIS class moves towards a unimodal shape with more than 80% of the CpGs having a methylation content between 0 and 0.2 (Fig. 3A). The few methylated CpGs are predominantly found at the 5’ end of this region class (Fig. 3B). Conversely, and in addition to the 2000-Down-TIS also the enhancer class exhibits a bimodal shape. Here, approximately two thirds of the CpGs are within the range of 0 to 0.2, one sixth between 0.2 to 0.8 and another sixth between 0.8 to 1 (Fig. 3A). The 2000-Down TIS region class shows a strong increase in methylation on CpGs from the 5’ to the 3’ end (Fig. 3B) -not surprising given it is, by construction, a cutout on the left edge of the gene. Enhancers are the only class of regions to exhibit a symmetrical pattern in terms of methylation percentage and CpG density (Fig. 3B).

**Table 1.** Average similarity of methylation fractions between and within species in percent.

|  | 2000-Up-TIS | 2000-Down-TIS | gene body | enhancer |
| --- | --- | --- | --- | --- |
| Interspecies – comparison of species' consensi |  |  |  |  |
| human-chimpanzee | 95.4 | 93.5 | 90.0 | 92.9 |
| human-gorilla | 94.9 | 92.7 | 87.9 | 92.0 |
| human-orangutan | 94.6 | 92.6 | 88.7 | 91.8 |
| chimpanzee-gorilla | 95.0 | 92.8 | 87.6 | 92.1 |
| chimpanzee-orangutan | 94.7 | 92.7 | 88.7 | 91.9 |
| gorilla-orangutan | 94.6 | 92.5 | 87.2 | 91.3 |

**Fig. 3.**
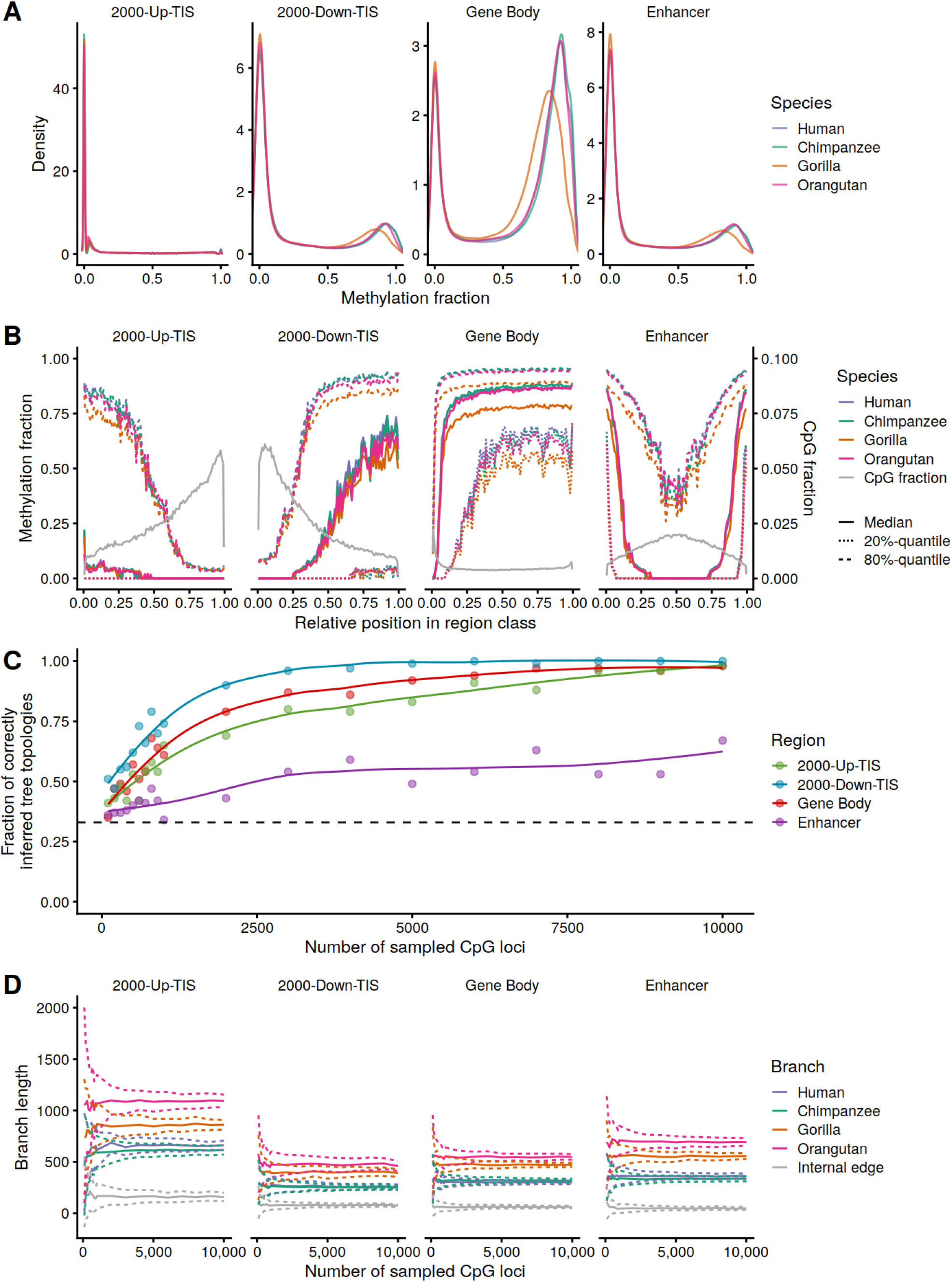
A) Distributions of methylation fractions in the examined regions and species. B) Methylation and CpG fractions by relative position in region class. C) Comparison of examined regions by the fraction of correctly inferred tree topologies. The total number of reconstructed trees per fixed amount of input data (i.e., Number of CpG loci) was always 100. D) Dispersion of branch lengths in the examined regions. The dashed line indicates the probability of randomly reconstructing the correct tree topology (one third). A,B,D) The given numbers of CpG loci were drawn from the respective region-wise methylation fraction alignments with the following total numbers of evaluable CpG loci: Up-TIS-2000 – 126,560; Down-TIS-2000 – 134,686; Gene body – 441,286; enhancer – 271,195.

Expectedly, the average similarity of the methylation fractions varies more between the regions under consideration than between species pairs – reflecting the distinct functional roles of the regions. Interspecies similarity ranges from 87.2-90.0% in the gene body to 94.6-95.4% in the 2000-Up-TIS region (Table 1).

To model a null-hypothesis for the analysis of region-specific similarities, we repeatedly uniformly (n=1000) drew random pairs of methylation rates from the class-specific background distributions and calculated expected differences. Based on this, we found that the level of region-specific similarity between species pairs is significantly higher than expected (p < 10^−3^, Table 1). These initial, descriptive results suggest that the methylation data contains evolutionarily conserved, phylogenetically analyzable signals. Despite a high degree of similarity probably dominated by region-specific biological functions, differences at this rather coarse grained level already reflect the phylogeny of the great apes.

### Tree reconstruction works best for TIS-downstream and gene body classes

To get a more detailed view, we compared the classes of regions in terms of their potential to reconstruct correct tree topology using the methylation data. With the default method, the 2000-Down-TIS region contains the highest phylogenetic signal (Fig. 3C). Notably, this class has by far the highest proportion of hemi-methylation (0.2-0.8). It seems plausible that CpGs, which are neither fully methylated nor demethylated across the tissue, also have the technical ability of being phylogenetically most informative. It is well documented that an increase in methylation downstream of the transcription start site and thus potentially overlapping with translated regions may have a strong suppressive effect on gene expression (confer Fig. 3 of (Ehrlich and Lacey 2013); (Appanah, et al. 2007)). At the same time, the region overlaps with proximal parts of the gene body where a hypermethylation is frequently associated with strong expression (Zemach, et al. 2010). In conjunction with the high level of global similarity established above, these results provide further evidence that specific phylogenetic information is embedded in this regulatorily relevant region. In addition to informative signals in regulatory regions, also inter-species expression differences may contribute to the surprisingly high rate of correct reconstructions. The observation that tree reconstruction based on the gene body works second best supports this notion.

With the notable exception of enhancers, the proportion of correctly reconstructed tree topologies converges clearly towards 1 in all region classes, depending on the amount of available data (Fig. 3C). The result is similar concerning the second benchmark, i.e. whether the determined branch lengths converge with increasing data volume. This is the case for all branches of the great apes in all regions studied (Fig. 3D).

### Methylation-based tree reconstruction may outcompete nucleotide-based reconstruction

For comparison, we juxtaposed methylation based reconstruction with classical nucleotide based reconstructions. For the gene body and Up-TIS-2000 classes, the fraction of correctly inferred trees is almost identical for methylation and nucleotide data. Most strikingly, using methylation data from the Down-TIS-2000 class reconstruction clearly outcompeted nucleotide-based reconstruction. Conversely, reconstruction based on methylation data obtained from enhancer regions essentially failed (Fig. 4A). In the light of high similarities of methylation fractions in this region (Table 1), this result is most surprising.

**Fig. 4.**
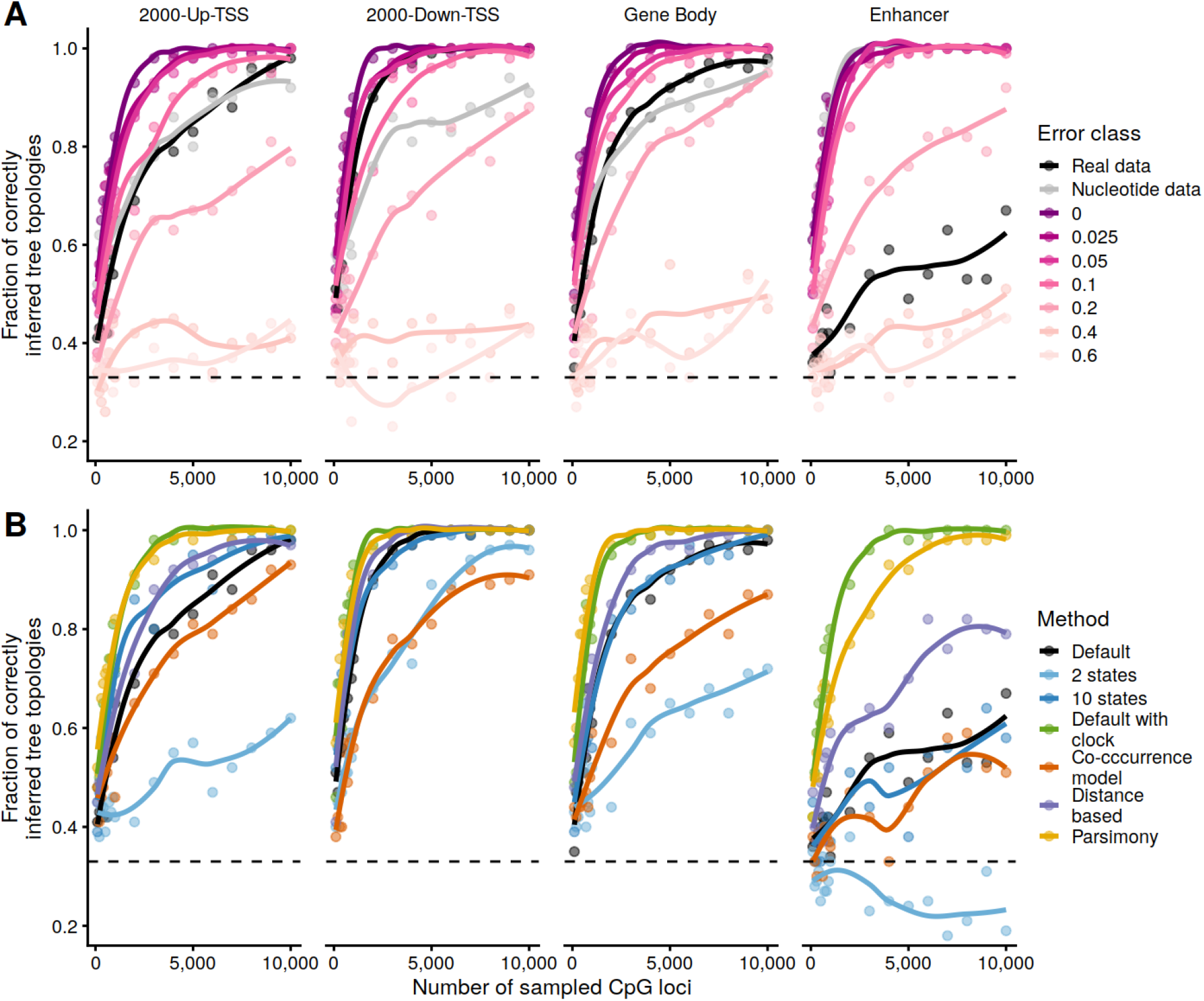
A) Model vs reality. The performance difference between the simulated data and the real data gives an estimate of how well the model matches reality. We first pretended that the model of the default method perfectly reflects reality: this model and the known phylogenetic tree of the great apes were used to generate artificial alignments. We then reconstructed phylogenetic trees from these alignments using the same model and applied the quality scale of fraction of correctly reconstructed tree topologies. To quantify the difference of model and reality, we additionally applied defined error terms, i.e. fractions of the standard deviation of the standard normal distribution, to the artificial alignments before tree reconstruction. B) Comparison of the different methods (or deviations of our standard method) by fractions of correctly reconstructed tree topologies. The given numbers of CpG loci were drawn from the respective region-wise methylation fraction alignments with the following total numbers of evaluable CpG loci: Up-TIS-2000 – 126,560; Down-TIS-2000 – 134,686; Gene body – 441,286; enhancer – 271,195. A-B) The total number of reconstructed trees per fixed amount of input data (i.e., Number of CpG loci) was always 100. The dashed line indicates the probability of randomly reconstructing the correct tree topology (one third).

To additionally quantify the performance of methylation based reconstruction, we generated artificial MFAs in complete analogy to nucleotide-or amino acid based alignments frequently used in the assessment of reconstruction algorithms or evolutionary models (e.g. (Rosenberg and Kumar 2001; Zhang, et al. 2005; Shavit Grievink, et al. 2010; Zaheri, et al. 2014). Using these MFAs, we reconstructed phylogenetic trees and determined the fraction of correct topologies (Fig. 4A). The procedure was repeated after adding different levels of artificial noise to the simulated methylation fraction alignments. At a noise level between 0.1 and 0.2 standard deviations the reconstruction performance of simulated data is at par with real data derived from gene body and Up-TIS-2000. Consistently, enhancers perform worse with values corresponding with noise between 0.2 and 0.4 standard deviations, while the Down-TIS-2000 class shows substantially better benchmarks with values between 0.05 and 0.1 standard deviations.

In summary, the reconstruction of phylogenetic trees from DNA methylation data seems to work well. The performance compared to nucleotide data is astonishing since methylation can be expected to be subject to frequent changes. Unlike DNA sequences, they are influenced by circadian rhythms, environmental factors or diseases. Clearly, in practice, the success of a methylation based reconstruction critically depends on the amount of available data, e.g. the read coverages achieved in WGBS experiments, the quality of reference genomes and sequence alignments. Furthermore, the reconstruction of phylogenies in single genes is naturally limited by the number of available data points. While the body of a protein coding gene has often more than 10,000 nucleotides, the measurement of the methylome is restricted to only 200 CpGs (Fig. S8). Experimental or biological noise may thus easily lead to faulty reconstructions. Unlike the genome, which is almost identical in all cells of an individual, the epigenome may reflect tissue-specific phylogenetical information. Thus, inter-species comparisons yield the potential of gaining insights into the evolution of tissues as opposed to the entire organism. The incorporation of other epigenetic data, e.g. histone modifications, will be helpful to illuminate such processes.

### Failure of tree reconstruction at enhancers

Triggered by the surprisingly bad tree reconstruction with enhancer methylation data, we characterized all sites individually with respect to their phylogenetic information. Theoretically, enhancers may escape our evolutionary models because of their functional heterogeneity, the concrete effect of enhancer-methylation on gene expression and their plasticity in terms of tissue-specificity or circadian effects (Angeloni and Bogdanovic 2019). Specifically, we assigned a score to each site quantifying its support of the correct tree topology and analyzed the score distributions. Compared to the most similar down-TIS-2000 class (cf. Table 1; Fig. 3A), the enhancers clearly contain more sites with negative information scores. Replacement of the least informative 10% of enhancer CpGs with the bottom Down-TIS-2000 CpGs (see Methods, Fig. S9), confirms the disproportionate impact of these sites on tree reconstruction. Un-informative CpG-sites show a significantly higher methylation fraction than the average enhancer class site (almost 0.6 vs. 0.4, p<2.2*10^−16^, Wilcoxon test). Furthermore, these sites are slightly enriched at 5’ and 3’ ends of enhancers (Fig. S10). The analysis of genes significantly affected by un-informative signals (FDR<0.1, Fisher-Test, 1092 out of 6328 significant, Table S2) reveals an enrichment of the IL12 pathway (BIOCARTA_IL12_PATHWAY, FDR = 0.03; BIOCARTA_NO2IL12_PATHWAY, FDR = 0.05) as well as leukocyte and lymphocyte differentiation (GO_LEUKOCYTE_DIFFERENTIATION, FDR = 0.01; GO_LYMPHOCYTE_ACTIVATION, FDR = 0.09). Since immunity-related genes themselves are targets of rapid evolutionary changes (Shultz and Sackton 2019), it is tempting to speculate that also sudden methylation changes within associated regulatory elements contribute to shaping the evolution of the immune response. Our data supports this hypothesis, as CpG methylation would be phylogenetically informative only at very short evolutionary distances.

### Non-binary models of CpG methylation work best for tree reconstruction

As shown in Fig. 4B, the model selection has a critical impact on phylogenetic reconstructions based on methylation data. The default No Jump Model is based on the idea that within short periods of time it is only possible to switch to adjacent states (Fig. 2). If a change from state A to a non-adjacent state B is to be made over a longer period of time, all intermediate states between A and B must be transited. With the alternative Co-occurrence Model, we permit sudden changes to non-adjacent states. Interestingly, the Co-occurrence Model consistently performs much worse than the No Jump Model across all regions. This behavior immediately raises the question to which extent such states are also functionally relevant. In contrast, when increasing the number of states to 10 typically no or only minimal improvement is observed. Naturally, the precise resolution of tissue-wide methylation fractions critically depends on the read coverage. Thus, for the data used in this study, a ten-state model likely is too fine-grained and over-specified.

### Tree reconstruction from DNA methylation data faces similar challenges as for nucleotide data

The presented default method builds on assumptions frequently made in the analysis of evolutionary relationships. To investigate the influence of those assumptions, we added a model with a molecular clock (1), used Neighbor Joining as a distance-based method for tree reconstruction (2), and applied the Maximum Parsimony principle (3; Fig. 2). All of the three models consistently perform better than the default method across all regions as measured by the proportion of correctly reconstructed topologies (Fig. 4B). For several reasons, it is frequently assumed that Maximum Likelihood is preferable to Maximum Parsimony or distance-based methods like Neighbor Joining (Felsenstein 1978; Hasegawa, et al. 1991; Kuhner and Felsenstein 1994; Guindon and Gascuel 2003). On the other hand, simulation and empirical studies at the nucleotide level have demonstrated that the fraction of correctly reconstructed topologies is often smaller as compared to other methods (Kolaczkowski and Thornton 2004; Yoshida and Nei 2016). Since the phylogenetic reconstruction for great apes clearly benefits from the molecular clock assumption on the nucleotide level (Moorjani, et al. 2016; Besenbacher, et al. 2019) and that the evolutionary rates at CpG dinucleotides are the most similar (Kim, et al. 2006), it can be expected that the molecular clock also improves the reconstruction performance based on MFAs. However, similar to the situation on the nucleotide level, this observation may not be generalized. Therefore, we have investigated the performance of the three methods in a scenario in which the evolutionary rates of the species diverge more strongly. Specifically, we carried out the simulations described above with a substantially higher evolutionary rate for one species than for the others (Fig. S11). Similar to the situation on the nucleotide level, maximum parsimony, and models that assume a molecular clock fail under this condition (Felsenstein 1978; Bergsten 2005). Also, the Neighbor Joining method performs significantly worse than our default method in these scenarios (Fig. 5A).

**Fig. 5.**
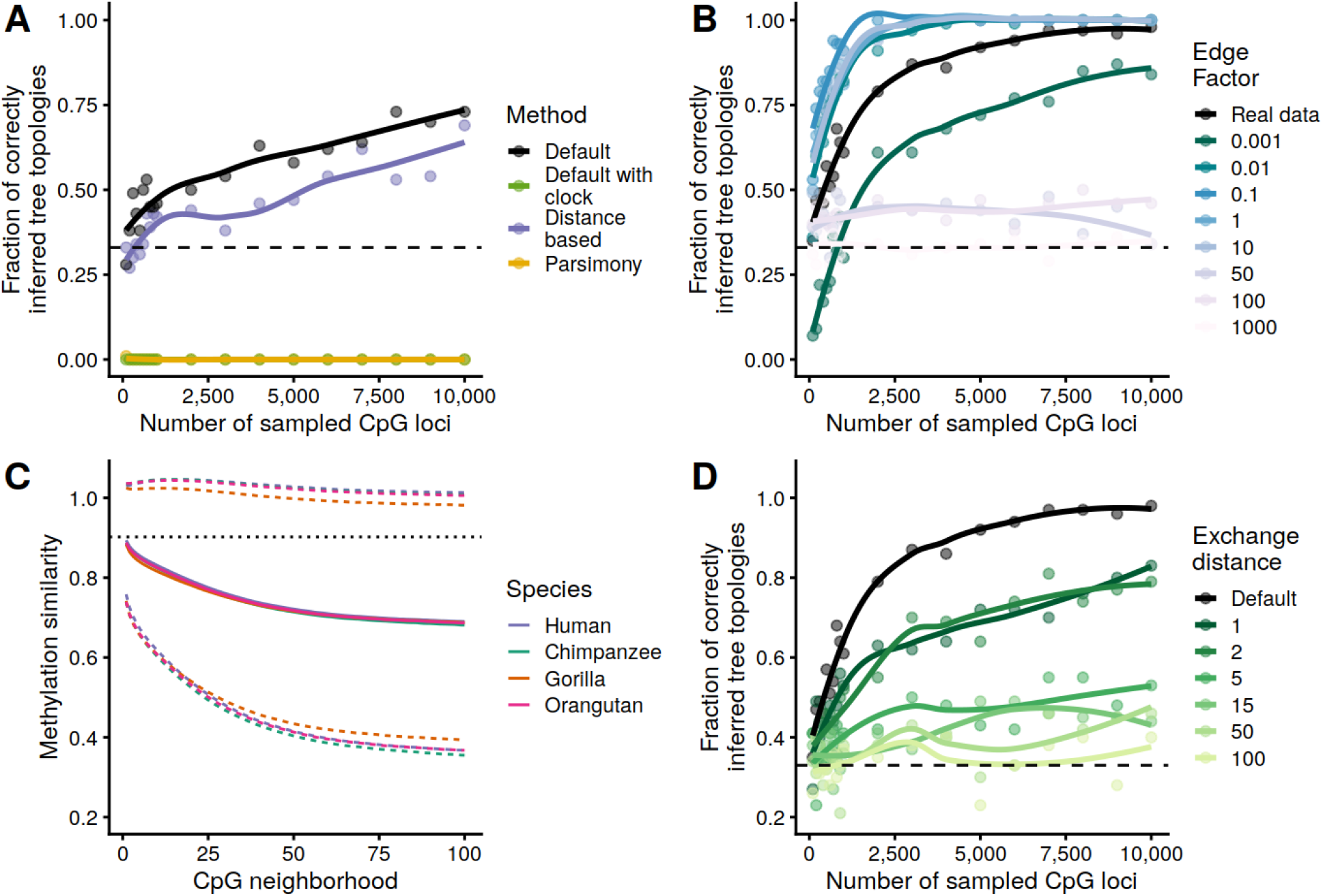
A) Comparison of selected methods in a long-branch attraction scenario. We generated artificial alignments using the model of our default method and a modified version of the great apes’ phylogenetic tree. The tree used for alignment generation was modified in such a way that one terminal branch was given a multiple of its original length. Then, trees were reconstructed from the alignments and the scale of the fraction of correctly inferred topologies applied. B) Resolution limits depending on the branch length. The same procedure as in A) was followed. The known phylogenetic tree of the great apes was used, however, for alignment generation, whereby all branch lengths were multiplied by fixed factors (Edge factor). C) Methylation similarity as a function of CpG neighborhood. We define the x-neighborhood of a CpG as the set consisting of the x-nearest CpG upstream and the x-nearest CpG downstream. Shown are the average similarities (solid line) and standard deviations (dashed line) of the methylation fractions for all corresponding x-neighborhood pairs. For comparison, the average similarity of the orthologous human-chimpanzee methylation fractions is also shown (dotted line, see also Table 1). D) Local signal resolution/alignment sensitivity. Phylogenetic trees were reconstructed from the real data set (great apes) using modified methylation fractions alignments. The alignments were modified so that methylation fractions were exchanged with a certain probability for a random methylation fraction of the same species within the given exchange distance. A, B, D) The dashed line indicates the probability of randomly reconstructing the correct tree topology (one third). A-D) The total number of reconstructed trees per fixed amount of input data (Number of CpG loci) was always 100. The region examined here was the gene body. The total number of evaluable CpG loci in this region is 441,286.

Based on the above-justified assumption that the default method sufficiently recovers the “true” phylogeny of species, we tried to obtain a rough estimate about the evolutionary distances to which it can be applied. For this purpose, we carried out a benchmarking where the branch lengths of the trees used for simulation were multiplied by a fixed factor. Our results indicate, that the default method should work similarly well for evolutionary distances ranging from one percent to 10 times the real data scenario (great apes) (Fig. 5B). Given the evolutionary rates of the great apes’ example and assuming that their last common ancestor lived about 15 million years ago (Locke, et al. 2011), one can cautiously estimate that the method would be suitable for distances of several 100 thousands to more than 100 million years.

### DNA methylation is conserved at the individual CpG site level

To investigate the degree of local conservation of methylation fractions, we determined the similarity of methylation for neighboring CpGs, defined as 1 − |*X*_*i,a*_ − *X*_*j,a*_|, where *X*_*i,a*_ and *X*_*j,b*_ are the methylation fractions at CpG-site *i* and *j* in species *a*, respectively. This similarity is contrasted with similarities of fractions in the spacial neighborhood of each individual species. Specifically, we define the 1-neighborhood of a CpG as the set of the next upstream CpG upstream and the next downstream CpG (without the CpG in the middle). The 2-neighborhood consists of the next but one CpGs in both directions (without the three CpGs in the middle) and so on. Expectedly, the average similarity of methylation decreases with growing neighborhoods towards a baseline level in all classes of regions. In the 2000-Down-TIS region, for instance, a similarity level of about 93% in the 1-neighborhood, decreases to about 80% in the 25-neighborhood and remains almost constant from then on (Fig. 5C, Fig. S12A). Strikingly, the class-dependent relationship between neighborhood and methylation similarities is practically identical in all of the investigated species. To put this observation into context, we contrasted the averaged neighborhood similarities with the similarity of methylation rates between two species at a given CpG, i.e. the in-between similarity. In any given class, the in-between similarity is at least as strong as the 3-neighborhood across all species. For the species pair human-chimpanzee, for instance, the in-between similarity is greater than that of the 1-neighborhood. In other words, our data indicates that, although separated by millions of years of evolution, the methylation rate of a CpG in humans is on average more similar to its orthologous CpG in chimpanzees than to that of its closest neighboring CpG. (Fig. 5C, Fig. S12A). To further investigate this, we have randomly swapped the methylation fractions of the real ape data set within defined exchange distances before reconstructing the phylogenetic trees. Even an exchange distance of one results in a significant decrease in the proportion of correct topologies, which continues to increase as the exchange distance is enlarged (Fig. 5D, Fig. S12B). This clearly indicates that the methylation on the DNA strand is conserved with high local resolution. On the flipside, this result also underscores the dependency of such methylation based studies on high-quality alignments – very much similar to studies based on the nucleotide or codon level (Jordan and Goldman 2012).

### Transcription factors preferentially bind to epigenetically conserved gene loci

Based on the observations of high inter-species similarity of CpG methylation (Table 1, Fig. 3A, 3B) and successful tree reconstruction using these data (Fig. 3C, 4A), we calculated a gene-wise estimates of the evolutionary rates to illustrate potential benefits of studies into epigenomic conservation. As described above, studies on a single gene level critically depend on the amount of evaluable data. With the data set at hand, the criteria for a thorough single-gene-based hypothesis testing are limited. In the context of this work we restrict ourselves to representative examples. *CASTOR1* codes for a component of the MTORC pathways (Saxton, et al. 2016) and is an example of a modestly conserved gene according to our epigenomic measure (Fig. 6A), i.e. the estimated rate of evolution is almost exactly that of the region average. Upon closer inspection, however, we observe that the local methylation level is anticorrelated with the transcription factor binding density, i.e. the number of different binding transcription factors normalized by gene length, which we determined based on publicly available data (Karolchik, et al. 2004). We also found that the rate at which a genes’ methylation level evolves is globally anticorrelated with the binding density of the transcription factors (ρ = -0.22, p < 2.2*10^−16^, Table S3). As a second intuitive measure for assessing functional aspects of epigenomic conservation, we determined how much the respective gene tree deviates from the average tree of the region class (see Supplement methods for details). *C16orf86* codes for a probably functional but so far uncharacterized protein and an example of a strong deviation (Fig. 6B). According to the described measure and our used filter criterions, the gene shows the highest deviation from the expected tree. According to our model, this deviation can be attributed almost exclusively to an increased evolutionary speed in humans. Interestingly, however, methylation levels in humans were only locally significantly different to other species between nucleotide positions 180 and 530. This sequence between the mentioned nucleotide positions is partly coding for a fully conserved domain of unknown function (pfam15762, DUF4691) and shows a relatively high TFBS density in humans. It is tempting to speculate that in the other great apes the relatively higher methylation may leads to a lower transcription factor binding density in this region.

**Fig. 6.**
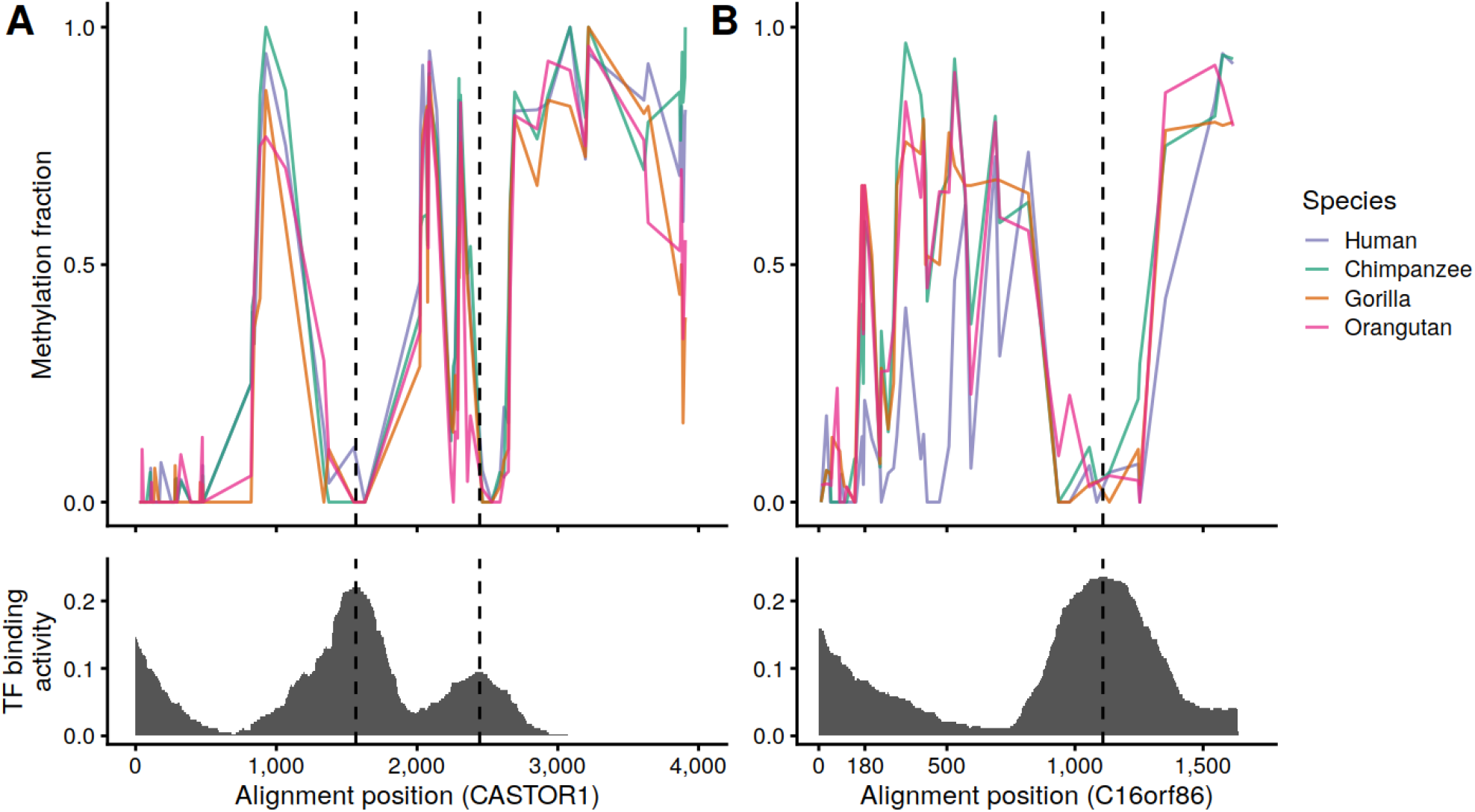
A) Measured methylation fractions and estimated local transcription factor binding density in the gene body of *CASTOR1*. The total number of evaluable CpG loci is 81. 19 CpG loci were skipped due to low read support (<10 reads in at least one species). B) Measured methylation fractions and estimated local transcription factor binding density in the gene body of *C16orf86*. The total number of evaluable CpG loci is 45. 4 CpG loci were skipped due to low read support (<10 reads in at least one species).

### Hints of accelerated epigenomic evolution for PRC2 protein binding sites

Based on our finding that transcription factor binding density appears to be negatively correlated with our measures for epigenomic evolution we were specifically interested to characterize individual transcription factors in terms of their evolutionary rates. To do this, methylation rates of aligned gene body CpGs overlapping with a specific TF were combined. In total, 297 out of 340 transcription factors comprised at least 1.000 CpG sites -the chosen minimum to include a TF in this analysis. The strongest deceleration on the human branch has been observed for RNA-binding-protein 34, RBM34. While little is known about the function of this gene, its binding to DNA might be influenced by the CpG-methylation.

Significantly accelerated rates (FDR <= 0.01; see Methods) were found for about a third of TFs (n=98) in humans (Fig. 7A). Strikingly, this set appears to enrich critical components of the polycomb repressive complex 2 (PRC2) complex, namely, YY1, EZH2, HDAC1, HDAC2, BMI1, and SUZ12 (BIOCARTA PRC2 pathway; FDR=0.092). Binding sites of the two histone deacetylases are also components of the significantly enriched telomerase pathway (Pathway Interaction Database; FDR = 0.0026) additionally comprising accelerated TFBS of XRCC5, MAX, SP1, IRF1, MYC, NBN, NR2F2, E2F1, SAP30, SIN3A and SIN3B. The strongest accelerations with a more than 50% increased evolutionary rate on methylation level were found for ZZZ3, CBX2, MYB, CDC5L, MIER1 and BRCA1. The zinc-finger ZZZ3, a component of the Ada-two-A-containing (ATAC) histone acetyl-transferase complex, has recently been described to function as a reader of histones regulating ATAC-dependent promoter histone H3K9 acetylation (Mi, et al. 2018). It is tempting to speculate that evolutionarily driven changes of ZZZ3 binding characteristics could have a decisive impact on species-specific gene expression. The chromobox homolog protein 2 (CBX2), also a reader of histone modifications, is a member of the polycomb repressive complex 1 (PRC1) (Vandamme, et al. 2011). It contributes to the repression of genes by binding to H3K9me3 and H3K27me3.

**Fig. 7.**
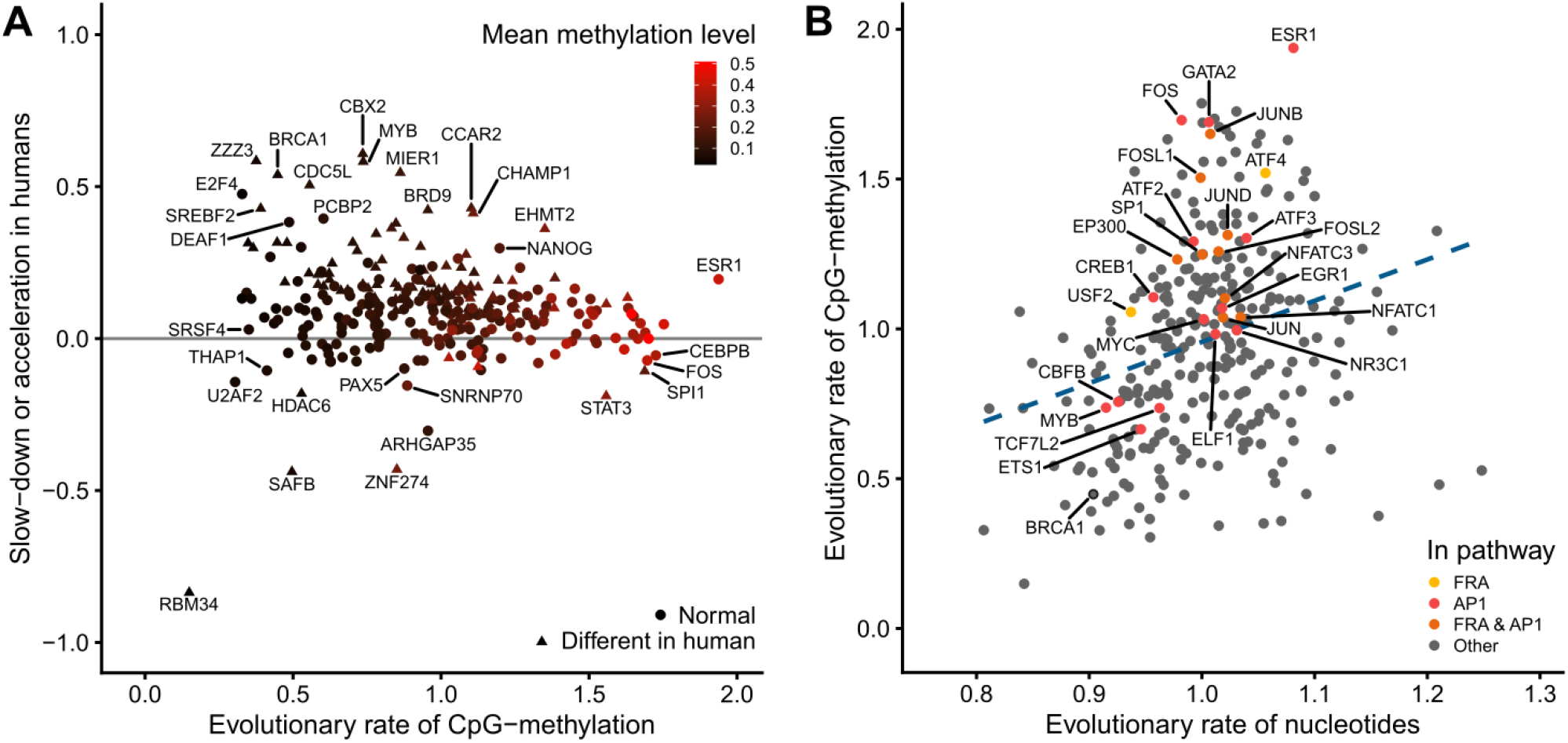
A) Estimated evolutionary rate of transcription factor binding sites on CpG-methylation level in great apes vs relative slow-down/acceleration of this rate in humans. Triangles mark transcription factor binding sites with statistically significant deviation of the human evolutionary rate (FDR<0.01, maximum likelihood ratio test, see *Methods* chapter *Methylation at transcription factor binding sites*). The color codes the mean methylation level across all evaluable CpG sites in the binding sites of the respective transcription factor. B) Estimated evolutionary rate of transcription factor binding sites on nucleotide vs CpG-methylation level. The dashed line indicates a simple linear regression of the two evolutionary rates. Color-coded are all names of transcription factors that are part of the pathways that are listed by the legend (Pathway Interaction Database (Schaefer, et al. 2009)). A, B) Shown are those 297 out of 340 transcription factors from the UCSC hg38 *Transcription Factor ChIP-seq Clusters* track (Karolchik, et al. 2004) that comprised at least 1.000 evaluable CpG sites.

One of the strongest evolutionary accelerations on the methylation level can be observed at the TFBS of BRCA1. Notably, the gene itself has been found to undergo a rapid evolution driven by positive selection altering its amino-acid composition as well its non-coding parts (Pavlicek, et al. 2004; Lou, et al. 2014). Involved in double strand break repair, BRCA1 is frequently mutated in hereditary forms of human breast cancers. Thus, our data suggests that the accelerated evolution of BRCA1 in humans compared to other great apes (Fig. 7A) has a measurable impact on the methylation landscape around its binding sites. At the same time, on the sequence level, BRCA1 binding sites evolution appears to be substantially decelerated (Fig. 7B). This marked lack of co-variance potentially supports the functional effects of changes in the BRCA1 gene itself. Notably, BRCA1’s direct interaction partner, the estrogen receptor ESR1, exhibits a comparably high evolutionary rate across all primates on the methylation level as compared to the sequence of its binding sites (Fig. 7A, 7B). The systematic comparison of sequence-based and methylation-based evolutionary rates reveals an acceleration of binding sites for transcription factors involved in neuron differentiation (GO_NEURON_DIFFERENTIATION, FDR = 0.03) and two highly connected pathways, AP-1 (FDR = 0.03, PID_AP1_PATHWAY) and FRA (ii, FDR = 0.02, PID_FRA_PATHWAY), due to the constitutive components of the heterodimeric AP-1, including, e.g., FOS, the ATF family, GATA2 and JunB (Fig. 7B). The proteins belonging to the AP-1 family play a critical role in numerous cellular processes. Notably, in cooperation with other cell-type specific factors it is involved in the activation of cell-type specific enhancers (Madrigal and Alasoo 2018). Hence, this result may be explained by species-specific cell compositions, systematic environmental or nutritional differences. Nevertheless, it is tempting to speculate that the enrichment of these factors may point to combinatorial changes of AP-1 composition during evolution.

## Conclusions

Here, we have systematically transferred classical methods of phylogenetics used to analyse nucleotide and amino acid sequences, to the field of epigenomics. Using empirical data from great apes, we demonstrate that phylogenetic trees can be correctly reconstructed from methylation data based on the fundamental principles of maximum likelihood, parsimony, and distance-based approaches. The problems and challenges are similar in many respects to tree reconstruction from nucleotide data, e.g. that parsimony and distance based methods are prone to long branch attraction. Nevertheless, we found that all examined regions with the notable exception of enhancers contain enough of phylogenetic information on CpG-methylation level to outcompete reconstruction with nucleotide data. The ability of the enhancers to escape the evolutionary model can be attributed to a relatively small set of CpG sites. The enrichment of these sites in quickly evolving immune-related genes highlights the importance of the epigenome for short-term evolutionary changes. Based on the empirical data and simulations, we showed that methylation levels are conserved at single CpG resolution. The methylation fraction of a CpG can usually be better predicted from the methylation fraction of an orthologous CpG in a species separated by millions of years of evolution than even from the methylation fraction of the closest CpG in the same species. We also found evidence that epigenetic conservation is associated with enhanced transcription factor binding density. Evolutionary rates on the nucleotide level were, as expected, found to be highly correlated with those CpG methylation level across transcription factor binding sites. However, significant deviations from this general trend are observed for binding sites of transcription factors associated with neuron differentiation and components of the heterodimeric AP-1 evolving significantly faster on the methylation level. Multiple examples provide hints that epigenomic re-modellers themselves could be critical components in the evolution of the human lineage. Significantly elevated evolutionary rate on methylation level in humans as compared to other great apes were found at TFBS of BRCA1, CBX2, ZZZ3, MIER1 and MYB. On a global level, critical components of the polycomb repressive complex 2 and members of the telomerase pathway show an accelerated CpG methylation evolution in humans.

In the future, when data will be collected for different epigenetic marks across multiple tissues, these methods would be helpful to test for accelerated or slowed epigenetic evolution affecting individual genes. This promises new insights into the evolution of ontogenesis, mechanisms of epigenetic gene regulation and possibly the formation of phenotypes based on these mechanisms.

## Supplemental methods

### Read mapping and quantification of methylation at individual CpGs

The soft masked versions of the following reference genomes were downloaded from ensemble release 95: GRCh38 (human), Pan_tro_3.0 (chimpanzee), gorGor4 (gorilla), PPYG2 (orangutan). Only chromosome entries from the reference genomes were considered for further analysis. Reads from the test data set were adapter clipped using TrimGalore (https://github.com/FelixKrueger/TrimGalore, version 0.4.5_dev, (Martin 2011)) mapped against the respective reference genome using segemehl, version 0.3.4 (Hoffmann, et al. 2014) with the parameter “-F 2” for the bs-seq protocol that was also used in the original study (Hernando-Herraez, et al. 2015). Subsequently, only unique mapping reads were considered as indicated by the NH:i:1 tag and duplicons were removed using samtools markdup, version 1.10 (Li, et al. 2009) (see Table S1 for mapping and duplicon rates). Methylation fractions at individual sites were determined using the haarz program from the segemehl package. Only methylation fractions within a CpG-context were considered for further analysis.

### Phylogenetic information of methylation values

To better elucidate the impact of individual methylation values for the reconstruction of the correct tree topology, we consider the phylogenetic information score *PIS*^*i*^ of site *i*. Let *ℒ*_*topology*_ be the maximized likelihood of the complete regions class’ data (with *k* CpG sites) and the unrooted *topology*, i.e.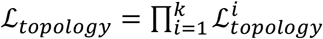. Then, we define *PIV*^*i*^ as follows:

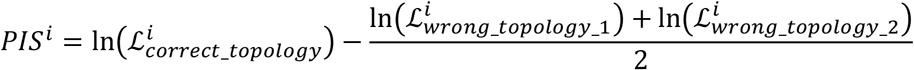

First, for each region class, sites are sorted with respect to their phylogenetic information. Subsequently, methylation values for sites within the lowest or highest quantiles are replaced with values from other classes. For instance, by removing 5% enhancer sites with lowest phylogenetic information with the same amount of sites sampled from the lowest 5% of Down-TIS-2000 class. Here, the performance of these mixed sets are evaluated by the correct tree topology benchmark (Fig. S9).

Specifically, we tested which enhancers were enriched for the 10% sites with lowest phylogenetic information score using the Fisher-test, a significance threshold of FDR < 0.1, and all examined enhancers as background. Corresponding interaction genes were counted as affected only if all their known enhancers showed an enrichment of those sites. Pathway/gene set enrichment was performed as described in *Methylation at transcription factor binding sites*.

### Methylation at transcription factor binding sites

To reveal potential functional consequences of epigenomic conservation, we focused on the methylation at known transcription factor binding sites (TFBS). For each of the 340 transcription factors, all CpG sites within the gene body class covering the binding sites of an individual *TF*, were combined into a methylation fraction alignment *X*^*TF*^. In addition, a methylation fraction alignment *X*^*TFs_combined*^ was created from the union of all CpG sites of the 340 transcription factors. To obtain a suitable reference, a phylogenetic tree was reconstructed from *X*^*TFs_combined*^ using the default method and the known great ape tree topology. Subsequently, for each individual transcription factor *TF*, the default method was used on *X*^*TF*^, forcing the branch lengths to be multiples of 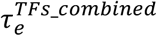 via the contraction/expansion factor 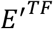

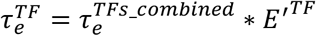

The evolutionary rate *E*^*TF*^ of *TF* was estimated as that instance of 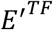 which resulted in the corresponding maximized likelihood 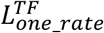 using the default method. To test for an acceleration/deceleration on the human branch, one additional methylation-based tree was reconstructed for each *TF* with the maximized likelihood 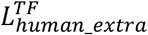. For this tree, all branches but the human branch were scaled with a fix contraction/expansion factor as before, while the human branch was free to vary. The actual phylogenetic hypothesis testing based on a likelihood ratio test then follows the standard procedure as described in the literature (see, e.g., (Huelsenbeck and Crandall 1997)) using 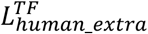 for the alternative hypothesis and 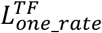 for the null hypothesis. Test results are expected to be chi-square distributed with one degree of freedom. We used the Benjamini-Hochberg method for multiple test correction (Benjamini and Y. 1995) and a significance threshold of FDR < 0.01 on transcription factor level (Table S3). For comparison, a nucleotide sequence-based estimate of the evolutionary rate at the TFBS was calculated with a co-occurrence model using the same multiple sequence alignments from which the corresponding MFAs were derived (see *Identification of orthologous defined regions* and *CpG Alignment*).

Subsequently, enrichment analyses were performed with the following gene sets taken from the Molecular Signature Database collections (MSigDB, (Liberzon, et al. 2015)): Biological Process Gene Ontology (The Gene Ontology 2019), Pathway Interaction Database (Schaefer, et al. 2009) and Biocarta pathway database. The analyses were restricted to gene sets containing at least 6 transcription factors and carried out using a fisher test followed by Benjamini-Hochberg correction as well as the above mentioned 340 transcription factors as background (Benjamini and Y. 1995). The significance threshold was set to FDR < 0.1. The test was applied separately for transcription factors with accelerated and decelerated evolution on the human branch. To identify pathways with significantly higher or lower evolutionary rates on methylation level as compared to the nucleotide level, we applied a t-test to the residuals of a linear model by contrasting residuals of the transcription factors in the respective pathway gene set vs. the residuals of the other transcription factors. Normality of the residuals was confirmed using the Shapiro-Wilk test.

Furthermore, we were interested in finding out whether individual genes regulated by a transcription factor themselves differ in terms of their epigenomic evolutionary rates. We reconstructed gene trees from methylation fraction alignments *X*^*g*^ of single genes using our default method and enforcing the correct great apes topology. We consider a single epigenetic evolution rate *E*^*g*^ of the gene *g* as well as the associated deviation *D*^*g*^ from an expected tree based on *E*^*g*^ and the average tree across all genes. Genes are sorted with respect to both metrics and correlated with the estimate of the transcription factor binding density *TFBD*^*g*^, i.e. the fraction of the 340 TFs having a binding site in the gene *g* divided by the gene length.

We define 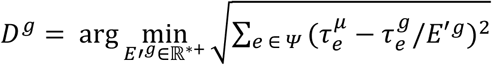 and *E*^*g*^ as that instance of *E*′^*g*^ that minimizes *D*^*G*^. In this context, *Ψ* is defined as the set of edges of the correct great apes tree topology, 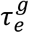 as the branch length of edge *e* of the tree reconstructed from *X*^*G*^, and 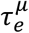 as the mean edge length of *e* from 100 reconstructions from as many drawings of *X* with each having a sample size of *N* = 10,000.

For correlations and examples, no genes were considered that reached either the minimum or the maximum of the allowed branch length of our optimization algorithm, which filtered out about 60% of the genes.

## Simulations

### Model vs. reality

We analyzed the capability of the default method to describe the real data. To this end, we repeatedly compared artificial scenarios reflecting the evolutionary methylation model with different levels of noise added to the real great ape methylation data.

First, we first converted the simulated states into simulated methylation fractions. Specifically, we drew a methylation fraction *M* for the respective state *s* from the interval corresponding to the discretization function *d* assuming a uniform distribution within the states. Subseqently, the noise *ε’* was drawn from a normal distribution with mean 0 and a fixed standard deviation *sd*. The new state *s*^*^ corresponding to *s* was then determined by

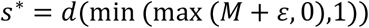

The entire approach described in this section was performed for different values of *sd*. The expectation is that the quality measures deteriorate with increasing *sd*.

In addition to the comparison with the performance of simulated data, we also compared the methylation data with the results on nucleotide data. Basis for these results were the same nucleotide alignments from which also our MFAs were derived (see *Identification of orthologous defined regions* and *CpG Alignment*). For this exercise, nucleotides were drawn from the whole alignment instead of methylation fraction from CpG sites. Since the No Jump Model underlying the default method was not applicable to the nominal scaled nucleotide data, the Co-occurrence Model was used.

### Resolution limits for reconstruction depending on the branch length

We extended our simulations to obtain an estimate of the evolutionary time scales on which the default method would be able to generate reliable results. For this purpose, we assumed the evolutionary rates estimated from data (great apes) to be fixed. Accordingly, the branch lengths of the tree used to generate the alignment were multiplied by fixed edge factors. The approach was performed with the following edge factors: 0.001, 0.01, 0.1, 10, 50, 100, 1000.

### Site-specificity of phylogenetic signals

To find out to what extent the phylogenetic signal is site specific, i.e. to answer the question of whether the evolutionary conservation rather affects single CpG sites or larger regions containing multiple sites, we resorted to the MFA’s obtained from real data (see *Preparation steps for empirical data*). Specifically, methylation fractions of randomly chosen sites were replaced with values from nearby CpGs of the same species.

Let 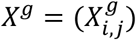, analogous to the definition of *X*, be the methylation fraction of the alignment of a gene *g* with *λ* homologous CpG sites; 1 ≤ *i* ≤ *λ*, 1 ≤ *j* ≤ *l*. For each alignment *X*^*g*^ we then generate a modified alignment *X*^*g*\*^ using a fixed modification probability *ρ* and a fixed modification distance *δ*

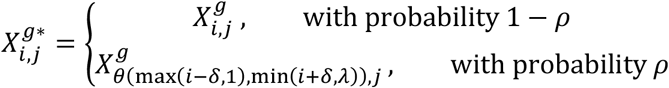

where *θ* is a function *θ*: ℕ^2^ → ℕ:

*θ*(*α, β*) = random number from {*α, α*+ 1,…, *β*}, with equal probability 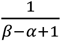 for all possible out*C*omes

with *α* ≤ *β*. By column-wise concatenation all *X*^*g*\*^ of a region, we create a merged modified alignment *X*^*^ analogous to *X*. Finally, we apply the same procedure to *X*^*^ as described earlier in *Evaluation strategy* and *Maximum likelihood* for *X*.

For this exercise, we chose a modification probability *ρ* = 0.7 and the modifying distance *δ*. The expectation is that the quality measures will deteriorate with larger values for *δ*.

### Long branch attraction

To evaluate the tree reconstruction methods with more divergent evolutionary rates than those of the great apes, we again created an artificial alignment of 100.000 sites, however, a phylogenetic tree that is more susceptible to long branch attraction instead of the great ape phylogeny was used (Fig. S11, (Bergsten 2005)) Finally, the procedure described in *Evaluation strategy* was applied with different tree reconstruction methods.

