## Supplement Figures S1-S12 for "The analysis of epigenomic evolution"

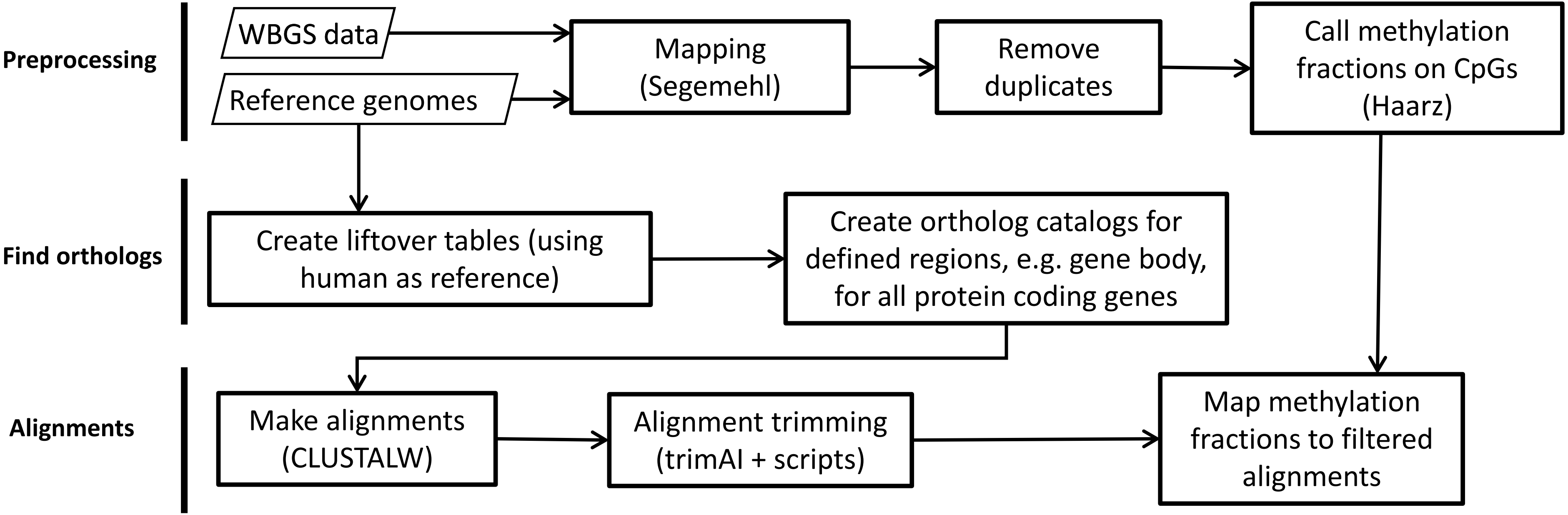


**Figure S1.** Workflow of preparation steps of the analysis. The subsequent steps are shown in Fig. 1B

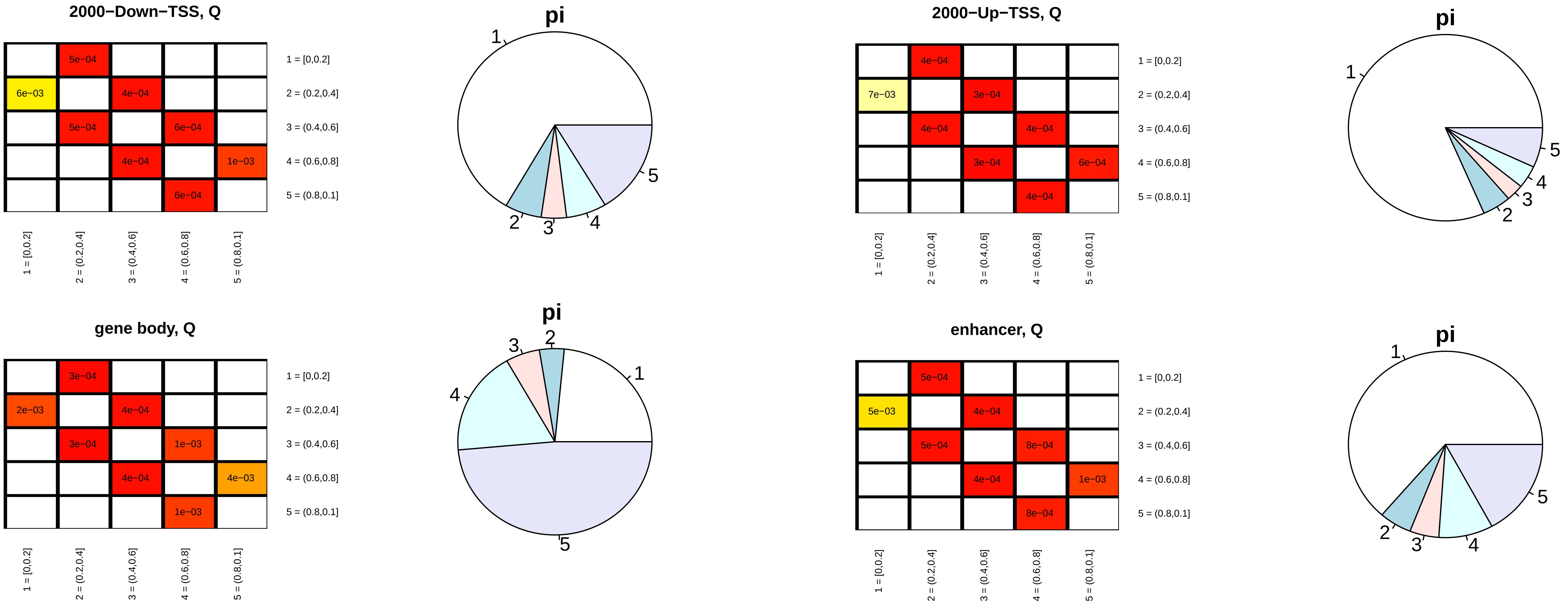


**Figure S2.** Parameterization of the No Jump Model for the four examined region classes.

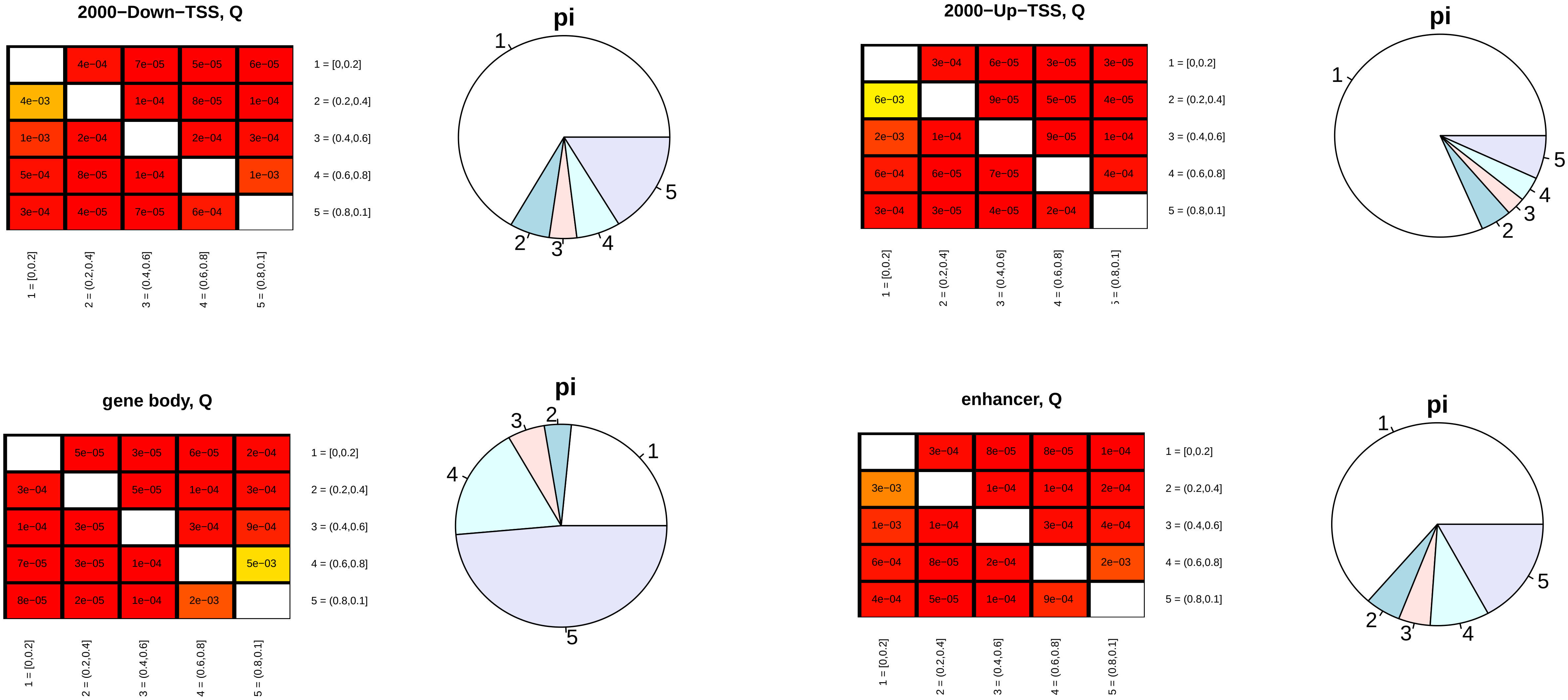


**Figure S3.** Parameterization of the Co-occurrence Model for the four examined region classes.

### Nominal scale (nucleotides etc.) → Fitch:

Assign set of letters *S(n)* for every node *n* in the tree:

- Initialization, for each leave *l*:

$$S(l) = \{observed\ character\ at\ l\}$$

- For each node *n* with children *x* and *y*:

$$S(n) = \begin{cases} S(x) \cap S(y), if\ S(x) \cap S(y) \neq \emptyset \\ S(x) \cup S(y), else \end{cases}$$

Example:

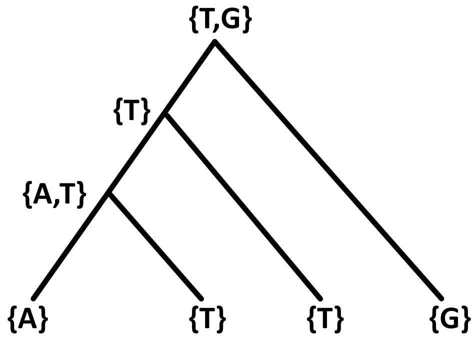


### Interval scale (e.g. methylation fraction):

Assign interval *I(n)* for every node *n* in the tree:

- Initialization, for each leave *l*:

$$I(l) = [observed\ number\ at\ l, observed\ number\ at\ l]$$

- For each node *n* with children *x* and *y*:

$$I(n) = \begin{cases} I(x) \cap I(y), if\ I(x) \cap I(y) \neq \emptyset \\ [\min(I(x)_{max}, I(y)_{max}), \max(I(x)_{min}, I(y)_{min})], else \end{cases}$$

Example:

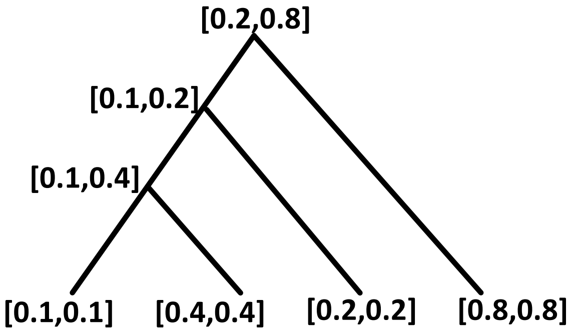


**Figure S4.** Comparison of nominal and interval scale pasimony algorithms: Bottom-up phase.

### Nominal scale (nucleotides etc.) → Fitch:

Assign letter *s(n)* for every node *n* in the tree:

- Initialization, for root *r*:

$$s(r) = arbitrary\ letter \in S(r)$$

- For each node *n* with parent node state *p*:

$$s(n) = \begin{cases} p, if\ p \in S(n) \\ arbitrary\ letter \in S(n), else \end{cases}$$

Example:

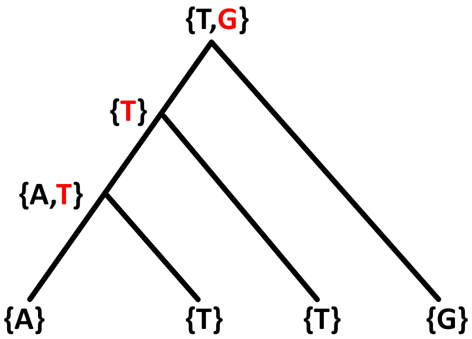


### Interval scale (e.g. methylation fraction):

Assign number *i(n)* for every node *n* in the tree:

- Initialization, for root *r*:

$$i(r) = arbitrary\ number \in I(r)$$

- For each node *n* with parent node state *p*:

$$i(n) = \begin{cases} p, if\ p \in I(n) \\ argmin_{x \in I(n)} |p - x|, else \end{cases}$$

Example:

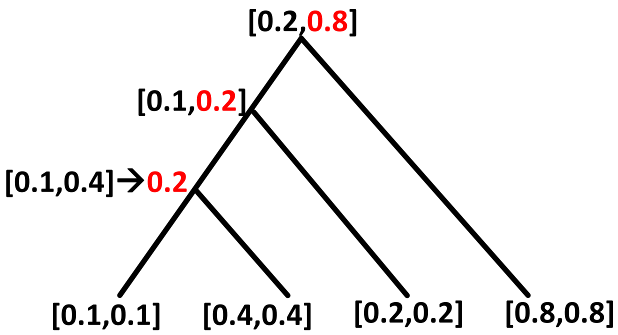


**Figure S5.** Comparison of nominal and interval scale pasimony algorithms: Top-down phase.

### Nominal scale (nucleotides etc.) → Fitch:

Assign costs *C(e)* for every edge $e = (n, m)$ in the tree:

- $C(e) = \begin{cases} 0, if\ s(n) = s(m) \\ 1, else \end{cases}$

Example:

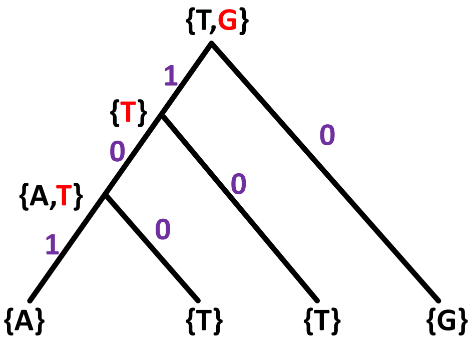


$$costs\ sum = 1 + 0 + 0 + 0 + 1 + 0 = 2$$

### Interval scale (e.g. methylation fraction):

Assign costs *c(e)* for every edge $e = (n, m)$ in the tree:

- $c(e) = |i(n) - i(m)|$

Example:

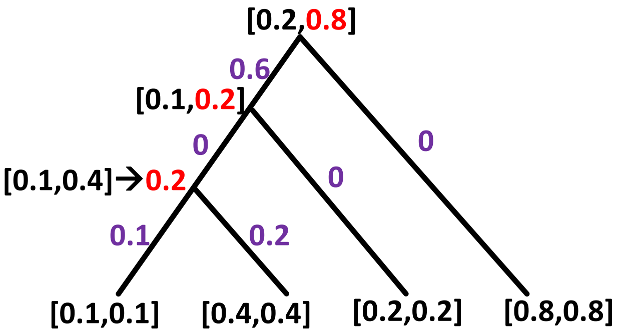


$$costs\ sum = 0.6 + 0 + 0 + 0 + 0.1 + 0.2 = 0.9$$

**Figure S6.** Comparison of nominal and interval scale pasimony algorithms: Cost determination phase.

**function** simulate_state($m$, $\tau_e$ ,$M$,$s$) {

**Input**: node $m = (\mathrm{T}, \eta)$ with list of lengths of outgoing edges $\mathrm{T}$ (empty for leaves) and corresponding list of child nodes $\eta$ (empty for leaves)

Edge length $\tau_e$ of incoming edge $e$ of $m$ (undefined for root)

model $M = (Q, \pi)$ with

$$\text{Rate Matrix } Q = \begin{pmatrix} Q_{1,1} & \cdots & Q_{1,n} \\ \vdots & \ddots & \vdots \\ Q_{n,1} & \cdots & Q_{n,n} \end{pmatrix}$$

Equilibrium frequency $\pi = (\pi_1, \ldots, \pi_n)$

state of parent node $s$ (undefined for root)

**Output**: list of simulated states for node $m$ and all descendant nodes

**if** $m$ is root **then** $\varphi := \pi$ //set probability vector for root to $\pi$

**else** {

$P := e^{Q * \tau_e}$

$\varphi :=$ s$^{\mathrm{th}}$ row of P

}

**declare** *simulated_states* as a list of states using nodes as element identifiers

*simulated_states*[$m$]:=draw a number from 1, …, n according to the probability vector $\varphi$

**if** length($\mathrm{T}$)>0 **then**

**for** $i$:=1 **to** length($\mathrm{T}$) *simulated_states*:=list_append(*simulated_states*, simulate_state($\eta_i, \mathrm{T}_i, M$, *simulated_states*[$m$]))

**return**(*simulated_states*)

}

**Figure S7.** Pseudo-code for recursive simulation of a state for each node of a phylogenetic tree, initiated given the root node $r$ and a Model $M$ consisting of a rate matrix $Q$ and an equilibrium frequency $\pi$: simulate_state($r$,undefined,$M$,undefined).

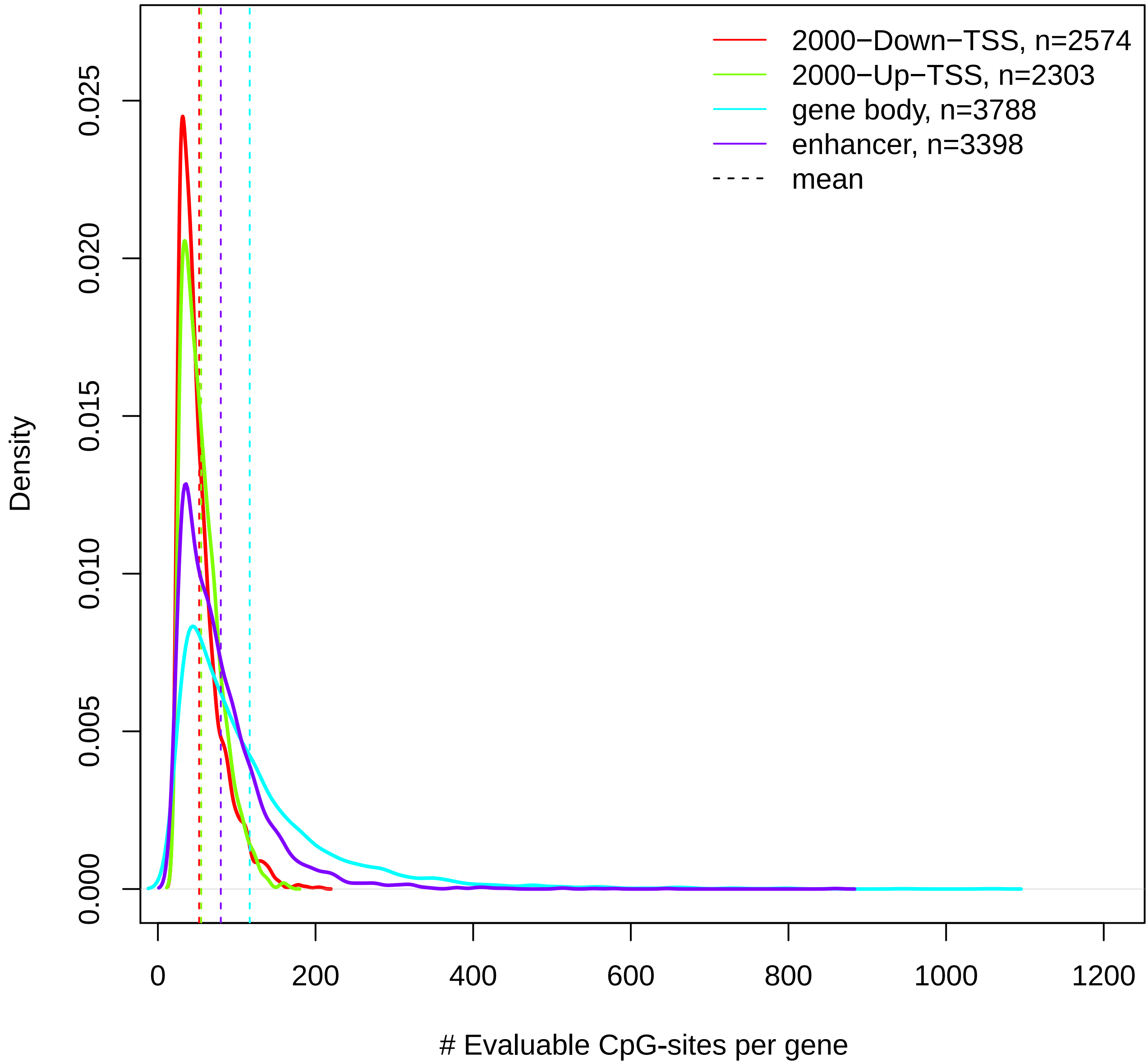


**Figure S8.** Distributions of number of evaluable CpG-sites per gene/enhancer. Only genes/ehnacers with at least 25 evaluable sites were included.

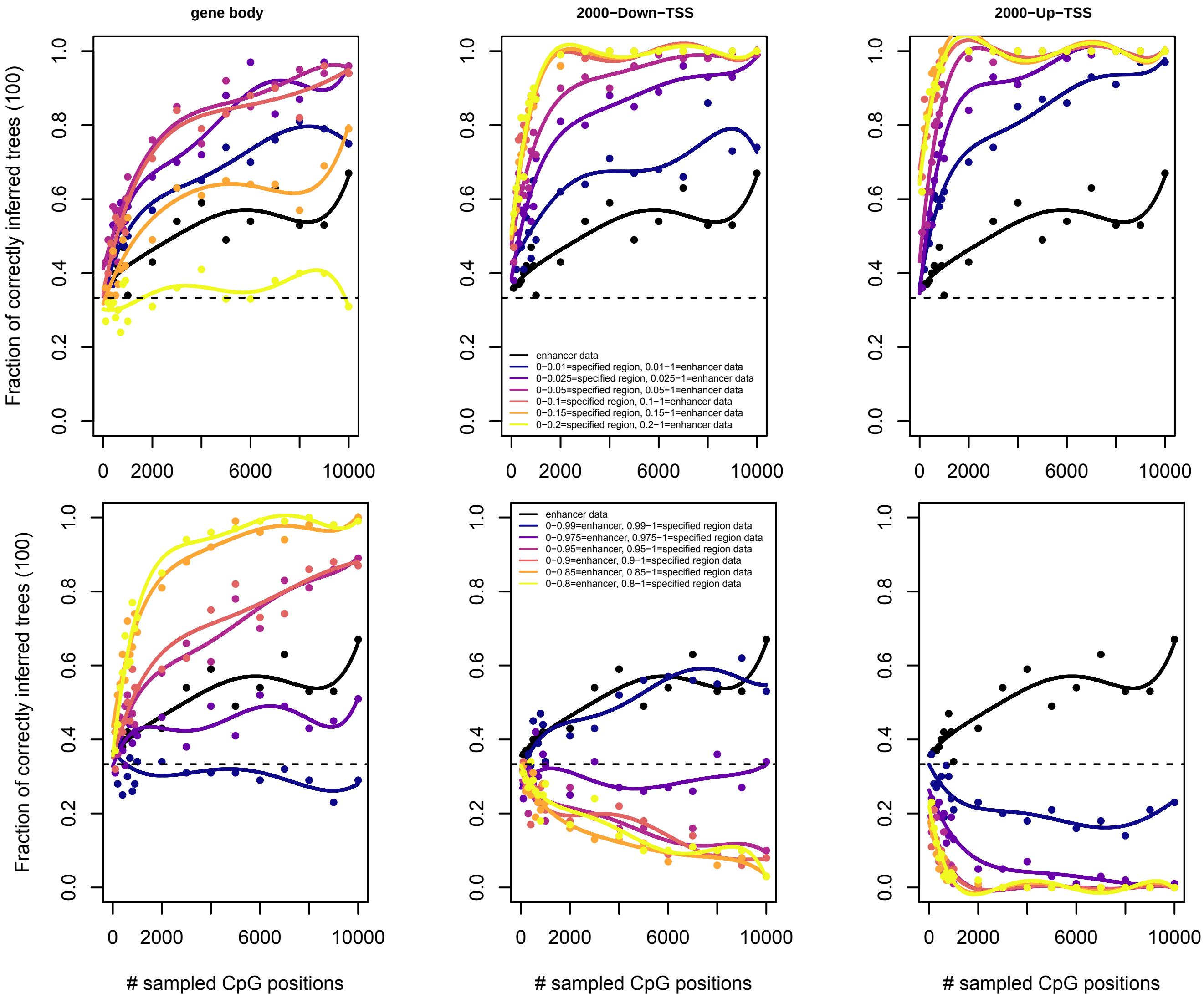


**Figure S9.** Performance of enhancer class data with sites of lowest (upper row) or highest (lower row) phylogenetic information value replaced by the respective sites of other region classes (columns).

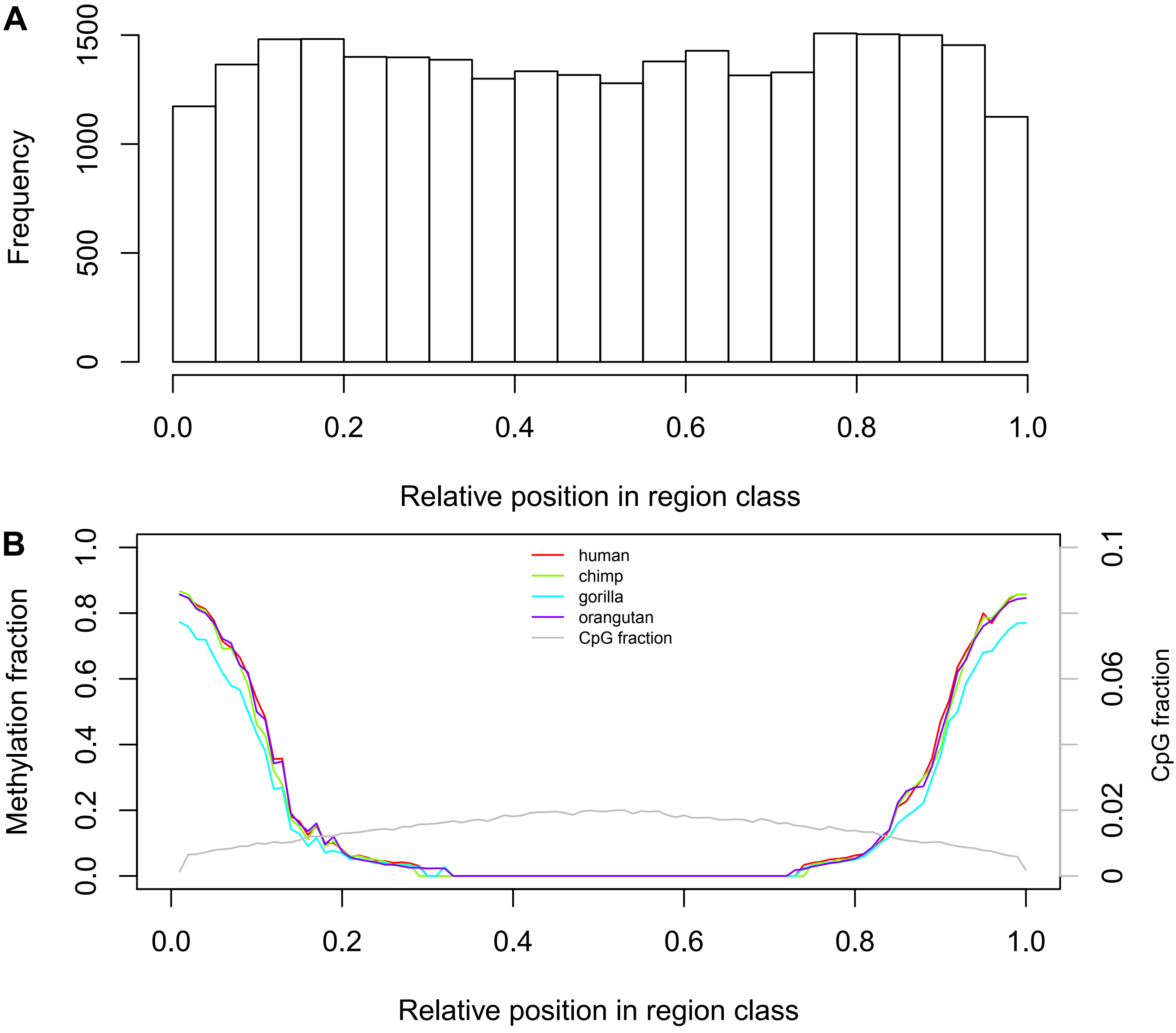


**Figure S10.** A) Regional distribution of 10% enhancer sites with lowest phylogenetic information value. B) For comparison, regional distribution of total enhancer sites (grey) and respective average methylation fractions.

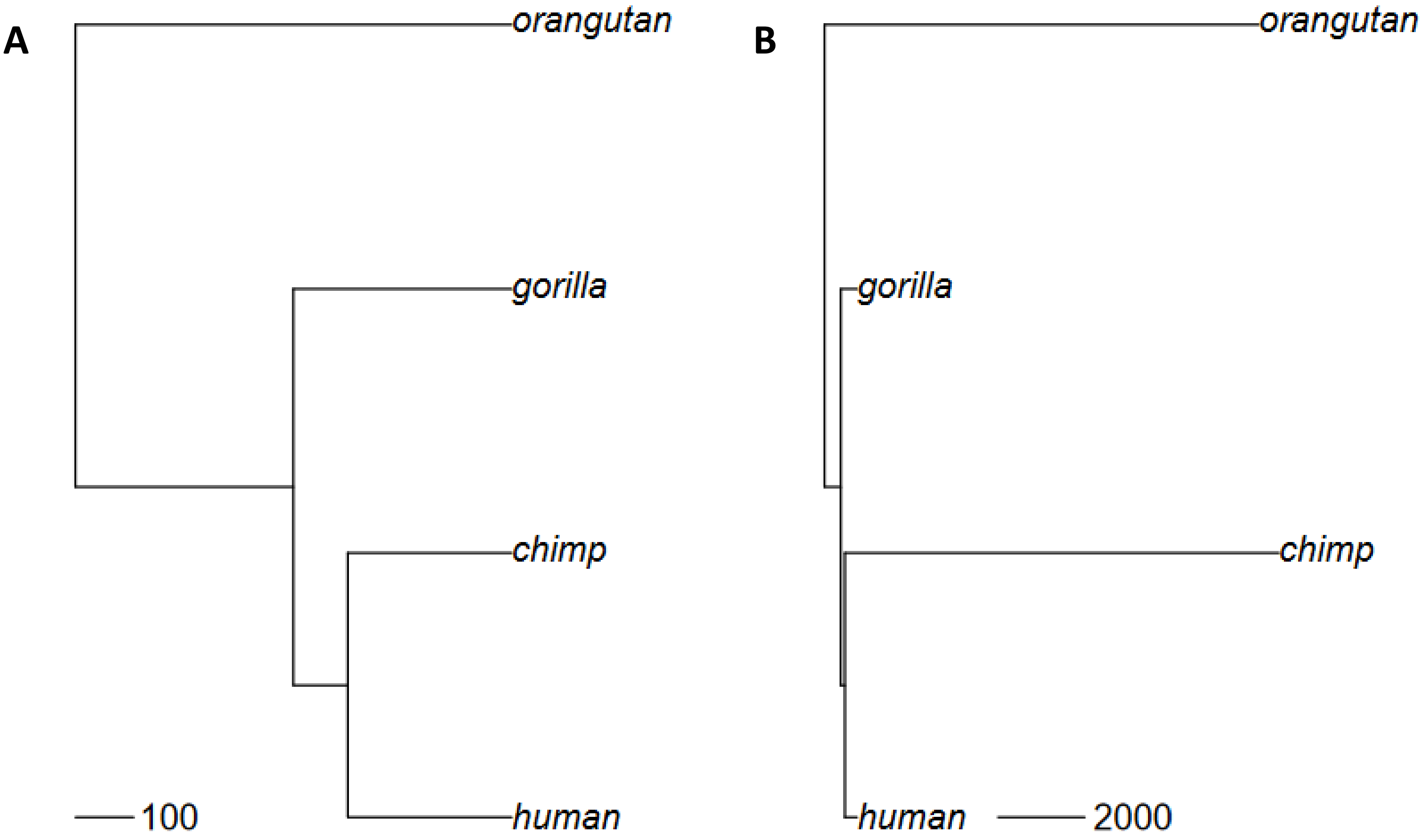


**Figure S11.** Phylogenetic trees used for simulations. A) Tree used for most analyses. Proportions of branch lengths were taken from Locke et al., Nature 469:529-533. B) tree used for long branch attraction scenario.

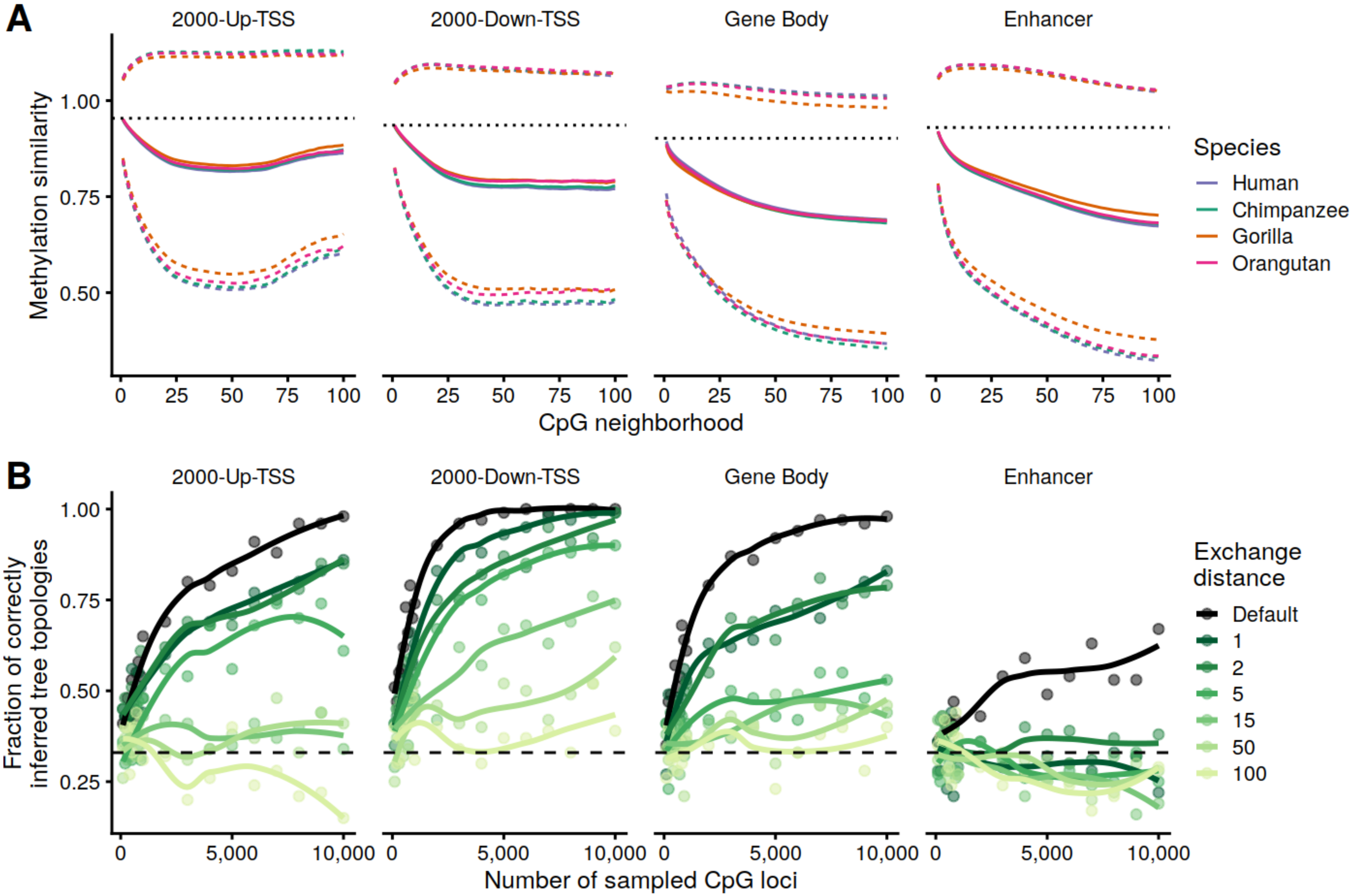


**Figure S12.** A) Methylation similarity as a function of CpG neighborhood. We define the x-neighborhood of a CpG as the set consisting of the x-nearest CpG upstream and the x-nearest CpG downstream. Shown are the average similarities (solid line) and standard deviations (dashed line) of the methylation fractions for all corresponding x-neighborhood pairs. For comparison, the average similarity of the orthologous human-chimpanzee methylation fractions is also shown (dotted line, see also Table 1). B) Local signal resolution/alignment sensitivity. Phylogenetic trees were reconstructed from the real data set (great apes) using modified methylation fractions alignments. The alignments were modified so that methylation fractions were exchanged with a certain probability for a random methylation fraction of the same species within the given exchange distance. The dashed line (at 1/3) indicates the probability of a randomly correct tree reconstruction.
